# Feeding your enemy’s enemy: Acidifying bacteria inhibit pathogenic bacteria more strongly with increasing glucose

**DOI:** 10.1101/2025.12.01.691202

**Authors:** Andrea R. Dos Santos, Clément Vulin, Sara Mitri

## Abstract

Classical microbiology has focused on directly suppressing pathogens using drugs, ignoring other harmless microbial species living alongside the pathogens. We now have a much better understanding of how species interact and affect one another’s growth within microbial communities, for example through chemical production. Here we capitalize on this understanding to demonstrate how one can manipulate and control the strength of interactions between bacterial species, and combine this with antibiotics to fully suppress and eliminate pathogens. Using experiments and a mathematical model, we first show how *Citrobacter freundii* can reduce the environmental pH to enhance the effect of ampicillin on the pathogen *Pseudomonas aeruginosa*. This negative interaction from *C. freundii* to *P. aeruginosa* can be strengthened by increasing glucose concentrations. Our proof-of-concept approach also worked against other pathogens: *Klebsiella pneumoniae* and *Agrobacterium tumefaciens*, and a different commensal: *Lactobacillus plantarum*, a common probiotic species. Overall, we show that taking advantage of the community and chemical context in which microbes live can help to develop efficient strategies to control them. In the medical context, this approach can help to eliminate pathogens thereby reducing our reliance on antibiotics.

## Introduction

Pathogens often share their environment inside the host with non-pathogenic microbes, also referred to as “commensals”. These different species can interact, influencing each other’s growth and survival positively or negatively depending on the species and the context. And yet, traditionally, pathogens have been studied in isolation, resulting in the discovery of many antimicrobial drugs that have become indispensable in treating microbial infections, and have saved millions of lives. But as we approach the crisis of increasing antimicrobial resistance globally (1), there is an urgent need to develop alternative, complementary strategies to effectively clear pathogens.

Thinking about antimicrobial administration in the presence of other microbes has a few important implications: antibiotics may cause collateral damage, killing or inhibiting untargeted competitors of the pathogen and making it easier for the pathogen to rebound (2–6). Non-pathogenic microbes may instead degrade the antimicrobials, thereby “cross-protecting” the pathogen (7), or enhance their effect to increase pathogen susceptibility to antimicrobials (8). The presence of other microbes can even impact resistance evolution, as shown in a simple cross-feeding community of *Escherichia coli* and *Salmonella enterica* (9).

Interactions between pathogens and non-pathogenic species need not just be an obstacle for eliminating pathogens or mitigating antimicrobial resistance evolution, though. Can one instead harness our understanding of these interspecies microbial interactions to develop new treatments with improved outcomes? In other words, can we control commensals to better compete with pathogens?

Competition between commensals and invading pathogens is an important component of pathogen resistance. For example, a diverse gut microbiome or certain members of the vaginal microbiome can prevent pathogen colonization (10). In this latter example, commensal species like *Lactobacilli* acidify the environment, reducing its pH to levels that are lower than the pathogen’s preference, thereby preventing it from invading (11–13). Such pH-mediated interactions are expected to be common, as all microbes are likely to modify their environmental pH (14, 15) and to respond to such modifications (16). Because each species’ metabolism and molecular machinery are adapted to function best within a specific pH range, pH modifications can potentially result in dramatic effects on fitness, by for example modifying the conformation of transport proteins in the bacterial membrane (17, 18). Because of this generality, environmental acidification (pH reduction) has parallels in other ecosystems, famously in food preservation, where fermentation by lactic acid bacteria leads to strong acidification (19).

Here we connect these ideas to test whether controlling the strength of negative interactions towards a pathogen through acidification can be used to complement the use of antibiotics. In previous work, we have shown that the sign and strength of interactions between two bacterial species can be controlled by modifying the concentration of a single chemical compound in the growth medium (20). In this study, we apply this same general approach to control the pH changes induced by one species to manipulate a pathogen and increase its susceptibility to antibiotic treatment. Using *Citrobacter freundii* to represent the acidifying commensal and *Pseudomonas aeruginosa* strain PA14 as the pathogen, we show that in the presence of *C. freundii, P. aeruginosa* is more strongly inhibited by the antibiotic ampicillin compared to when it is alone, and that this effect increases as we provide increasing concentrations of glucose to the co-culture, ultimately replacing the need for ampicillin altogether. A mathematical model allows us to identify key parameters to optimize *P. aeruginosa* inhibition and to predict commensal-pathogen species pairs for which this approach should work. We then experimentally test some of these pairs.

Our work links ideas on colonization resistance, food microbiology and interspecies interactions to demonstrate an effective strategy that can reduce the use of antibiotics. By taking advantage of the community context in which a pathogen (our enemy) is living, one can control it through the response of co-inhabiting species to resource concentrations (feeding the pathogen’s enemy), thereby increasing the strength of competition.

## Results

### Ampicillin can more strongly inhibit *P. aeruginosa* at low pH or in co-culture with *Citrobacter freundii*

We first sought to confirm previous findings that antibiotics – specifically ampicillin – can more strongly inhibit *P. aeruginosa* at an acidic pH (21–23). We performed a minimal inhibitory concentration (MIC) assay paired to a pH sensitivity assay where 10 ampicillin concentrations were combined with 6 pH values (Fig. 1A, S1). *P. aeruginosa* showed sensitivity to ampicillin (MIC = 200 *µ*g/mL at neutral pH, Fig. S2B) and to acidification (without ampicillin, growth is at 32% at pH 5 and no growth is observed below this value, n = 3). Gradually increasing acidification reduced the final yield of *P. aeruginosa* while ampicillin lengthened its lag phase (generalized linear model: pH *p<* 0.001 but not ampicillin *p* = 0.22 affected final yield, and ampicillin *p<* 0.001 but not pH *p* = 0.47 affected the lag phase, Fig. S1). Ampicillin and acidification together inhibited *P. aeruginosa*’s overall growth (generalized linear model: ampicillin *p<* 0.001 and pH *p* = 0.006 affected the AUC, Fig. 1A).

**Fig. 1.**
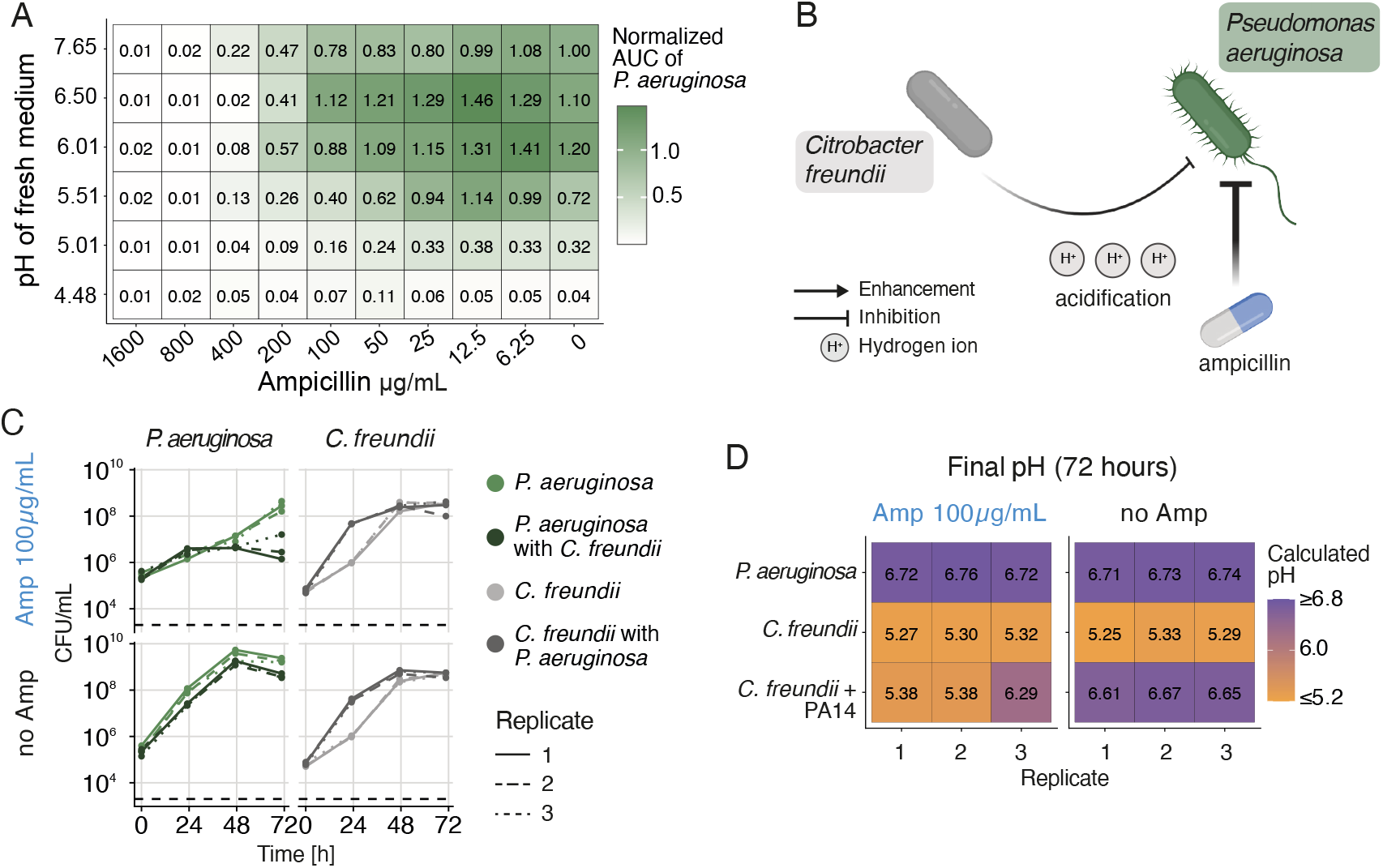
Acidification enhances the effect of ampicillin on the pathogen *Pseudomonas aeruginosa*. **(A)** Minimal inhibitory concentration (MIC) assay performed on *P. aeruginosa* using ampicillin, at 6 different initial pH levels: from 7.62 (fresh minimal medium, no pH modification) down to 3.62 (minimal medium + glucose 15 mM). The fresh medium was acidified using HCl. The OD_600_ was measured every 30 minutes over 72 hours and the area under the growth curves (AUC) calculated, normalized and plotted (n = 3). **(B)** Combining *Citrobacter freundii* with *P. aeruginosa* should enhance the effect of ampicillin on *P. aeruginosa. C. freundii* can significantly acidify the environment, which combined with ampicillin could more efficiently inhibit *P. aeruginosa*. **(C)** Growth of *P. aeruginosa* and *C. freundii* in monocultures and in co-culture, with or without ampicillin [100 *µ*g/mL] (medium = minimal medium + glucose 30 mM). Growth was determined by measuring CFU/mL over 72 hours (n = 3). The dashed line shows the detection limit. **(D)** pH from all conditions in panel C after 72 hours, measured using the absorbance of the pH indicator bromocresol purple at 588 nm (Purple above pH = 6.8 and yellow under pH = 5.2, see Methods).

Given that both low pH and ampicillin can negatively affect *P. aeruginosa*, we hypothesized that a species that lowers the pH should inhibit *P. aeruginosa* in the presence of ampicillin (Fig. 1B). To test this, we grew *P. aeruginosa* with *C. freundii*, a common host-associated bacterium. Upon growth alone in a glucose minimal medium, *C. freundii* acidified the medium from a pH of 7.62 down to ∼5.3 (Fig. 1C, D). Because we needed *C. freundii* to survive the ampicillin treatment long enough to acidify the environment, we isolated a *C. freundii* strain that was more resistant to ampicillin than *P. aeruginosa* (MIC 400 *µ*g/mL, Fig. S2A). To follow the pH of bacterial cultures over time, we used the non-toxic pH dye bromocresol purple whose absorbance at 588 nm can be correlated to the pH of the medium (purple in pH ≥ 6.8 and yellow ≤ 5.2, see Methods, Fig. S3).

When grown together without ampicillin, *P. aeruginosa* and *C. freundii* display weak interspecies interactions: compared to their growth in mono-culture, *P. aeruginosa* grows slightly worse with *C. freundii* (area under the growth curves AUC of mono-vs. co-culture *t*-test, p-value = 0.06), while *C. freundii* benefited from the presence of *P. aeruginosa* (AUC of mono-vs. co-culture *t*-test, p-value < 0.05). In co-culture with no antibiotics, acidification was in a similar range as when *C. freundii* was absent (Fig. 1D, right panel), suggesting that *P. aeruginosa* might cross-feed on the acidifying by-products produced by *C. freundii*, allowing the pH to remain close to neutral.

We next added ampicillin (100 *µ*g/mL) to the co-culture, and found that *C. freundii* could acidify the medium and reduce the population size of *P. aeruginosa* by approximately 100-fold in 2 out of 3 replicates (AUC of mono-vs. co-culture, t-test, p-value = 0.001). In the replicate where *P. aeruginosa* was inhibited the least, the final pH was also the highest (Fig. 1C and D, replicate 3, pH = 6.29), suggesting that *P. aeruginosa* inhibition depends on acidification. In summary, inhibition of *P. aeruginosa* by ampicillin can be increased if *C. freundii* can lower the pH.

### Model predicts that larger populations of *Citrobacter freundii* or more glucose can enhance the effect of ampicillin against *P. aeruginosa*

Before running longer-term experiments to explore whether we could eradicate *P. aeruginosa*, we sought to obtain more intuition on how the two-species system would respond to changes in experimental conditions. To this end, we built a consumer-resource model based on the dynamics displayed in Fig. 1, where the pathogen *B*_1_ (*P. aeruginosa*) and the commensal *B*_2_ (*C. freundii*) grow on a common resource *C* (glucose) in presence of an antibiotic (ampicillin). The commensal can produce an organic acid *A* while consuming *C* which acidifies the environment, acting synergistically with ampicillin to inhibit the growth of the pathogen. The pathogen consumes glucose as well as some of the produced organic acid. The parameters of the model were not fitted to the co-culture data, but were chosen to approximate our experimental observations. The model is described in detail in supplementary note S1.

Our overall goal is to eliminate *P. aeruginosa* at low ampicillin concentrations to reduce our reliance on antibiotics. To this end, we used the model to identify the conditions under which the pathogen would be excluded if we were to carry out serial transfers or growth-and-dilution cycles (Fig. 2A). We performed 2 parameter sweeps where we simultaneously varied the antibiotic concentration and either i) the initial population size of the commensal relative to the pathogen, or ii) the starting concentration of glucose (Fig. 2B). The model predicts that increasing either commensal population size or glucose concentration can lead to a quicker extinction of the pathogen, but only if antibiotics are used as well. If the commensal is absent, the pathogen survives till the end of the simulation under our chosen parameters even with antibiotics. If too little glucose is provided, neither species can grow enough to survive the 100-fold dilution. Overall, however, the model predicts that if the commensal is present, it should not be difficult to eliminate the pathogen in just a few transfers.

**Fig. 2.**
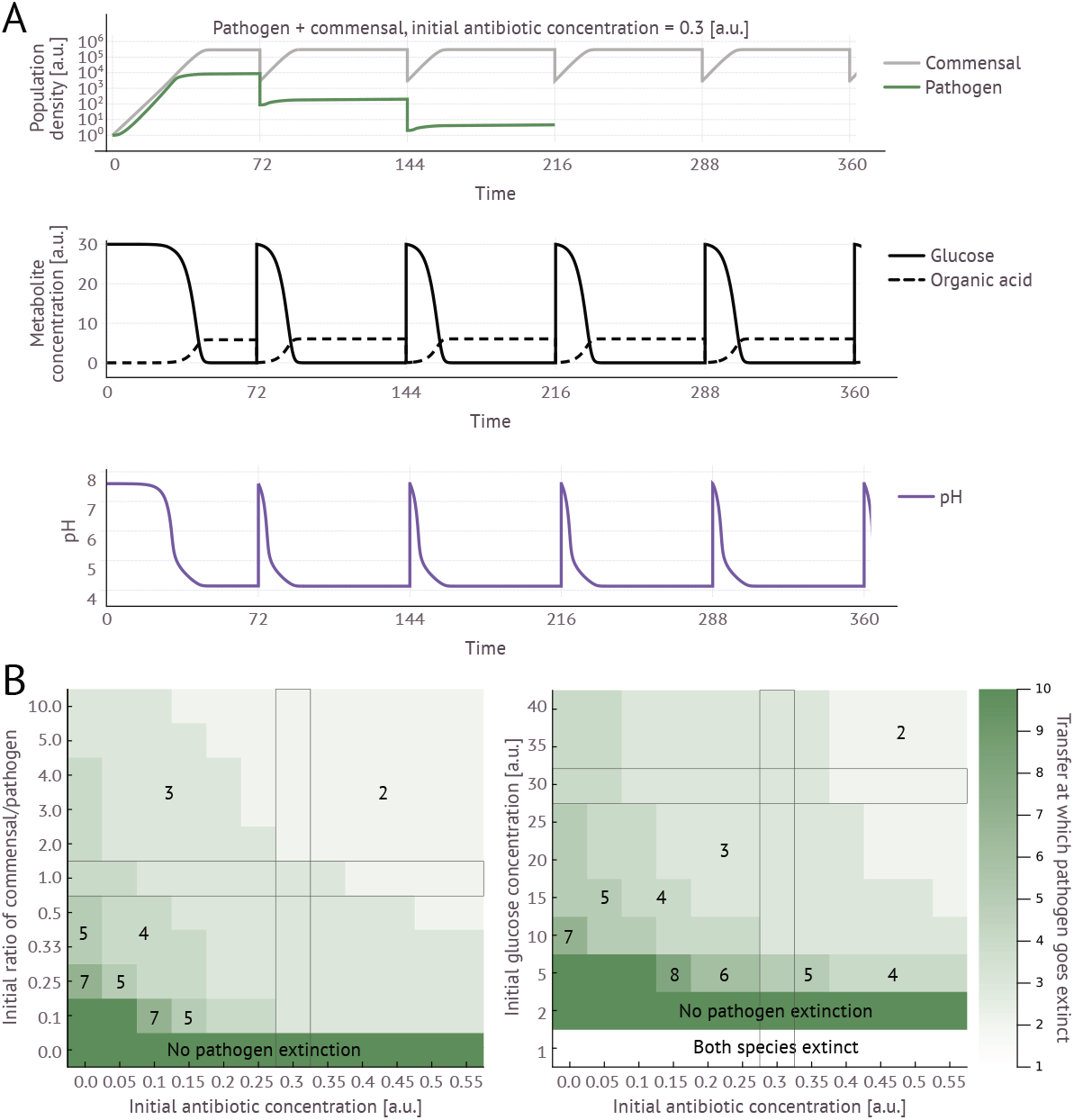
Mathematical model. **(A)** We simulate co-cultures of the pathogen (*P. aeruginosa*) and the commensal (*C. freundii*) in glucose in the presence of antibiotic (ampicillin) for 72 time steps (all arbitrary units, a.u.) followed by a 100× dilution into fresh glucose and antibiotic (top row, see Note S1). Growth and dilution cycles are repeated 10 times (only 5 are shown). As the species consume glucose, organic acids accumulate (central row), resulting in a drop in pH (bottom row). **(B)** We explore whether and how quickly the pathogen can be eliminated under different initial conditions. The pathogen is driven extinct more quickly when (left:) the commensal is initially more abundant or (right:) when glucose concentrations are higher, and when antibiotic concentrations are higher. Long rectangles show default parameter values (e.g. for simulations in panel A).

### Strong pH decrease by *C. freundii* at high glucose concentration eliminates *P. aeruginosa*

With our better understanding of the effects of population sizes, glucose and antibiotic concentrations, we next conducted the transfer experiments. We grew *P. aeruginosa* and *C. freundii* together and in isolation in different concentrations of glucose (15, 30 and 45 mM) and ampicillin (0, 50, 100 and 200 *µ*g/mL) with either approximately equal starting OD_600_ of *P. aeruginosa* and *C. freundii* (Fig. S4) or 10 × higher initial OD_600_ of *C. freundii* compared to *P. aeruginosa* (Fig. S5). Every 3 days for 15 days, we transferred 1% of the culture into fresh medium with the same initial concentrations of glucose and ampicillin. CFUs were quantified at the end of each transfer, and pH was followed over time by adding bromocresol purple to the growth medium (see Methods).

One of the clearest observations was that *P. aeruginosa*’s drop in population size significantly correlated with the final pH (Fig. S6, Pearson’s *ρ* = 0.78 or 0.94 for equal or different initial population sizes, respectively, *p* < 0.001 for both). Glucose concentration was an important determining factor: strong acidification (below 6) could only be achieved by *C. freundii* at glucose concentrations of 30mM or 45mM but not 15mM (Fig. S4A, B; S5A, B).

In partial contradiction with the predictions of the model, however, we found that initial population size of *C. freundii* had no detectable effect on *P. aeruginosa* extinction (generalized linear model with initial population size, antibiotic concentration, glucose concentration and partner species as factors, all were significant *p* < 0.001 except for initial population size *p* = 0.144). Unexpectedly, the concentration of ampicillin did not seem to affect the outcome much either, as final *P. aeruginosa* population size did not consistently correlate with ampicillin concentration (Fig. S6, Pearson’s *ρ* = −0.34 or 0.099, and p = 0.019 or 0.5 for equal or different initial population sizes, respectively). Indeed, some *P. aeruginosa* populations survived best at the highest concentration of antibiotics and did not always show increased resistance to ampicillin (Fig. S7). This led us to two hypotheses: (i) that the antibiotic concentrations being used (up to 200*µ*g/mL) were too low and (ii) that *P. aeruginosa* was forming biofilms protecting it from the antibiotic. The latter hypothesis was supported by visual examination of the wells (Fig. S8). In addition, in a shorter experiment where we mixed the cultures at regular time intervals to prevent biofilm formation, we observed that ampicillin was inhibiting *P. aeruginosa* as we had expected (Fig. S9, S10).

Encouraged by the effect of more frequent disturbance, we then repeated the growth-and-dilution experiment using higher concentrations of ampicillin (up to 800*µ*g/mL, above *P. aeruginosa*’s MIC) and by mixing the cultures twice per day using a pipette to prevent biofilm formation. As we observed very similar results at the two highest glucose concentrations (30mM and 45mM), we later carried out an additional transfer experiment using an initial concentration of 26mM of glucose (chosen based on a screen, Fig. S11) that showed an intermediate effect (Fig. 3).

**Fig. 3.**
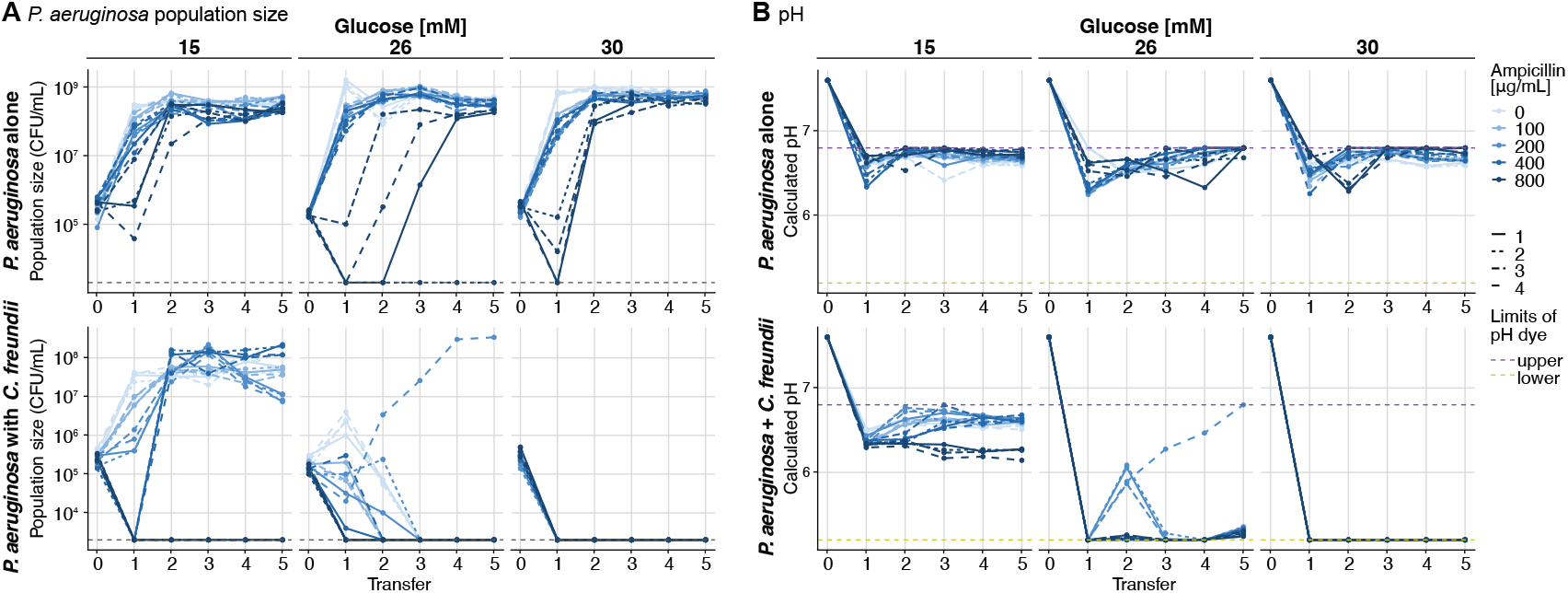
Experimental test of the effect of glucose and ampicillin on the inhibition of *P. aeruginosa*. **(A)** Population size of *P. aeruginosa* in various glucose and ampicillin concentrations when it is alone (top) or co-cultured with *C. freundii* (bottom) over 5 growth- and-dilution cycles (3 days of growth, 100X dilutions, or 1% transfers, n=4). For the bottom panel, the initial OD_600_ of *C. freundii* (0.01) is 10× higher than that of *P. aeruginosa* (0.001). Each sub-plot shows the population size of *P. aeruginosa* at the start of the experiment (transfer 0) and before each transfer. The corresponding data for *C. freundii* is shown in Fig. S12. The dashed grey line shows the detection limit. **(B)** pH calculated based on the absorbance at 588 nm of bromocresol purple (see Methods) at the start of the experiment and before each transfer when *P. aeruginosa* was alone (top) or with *C. freundii* (bottom). The pH of the fresh medium at the initial timepoint was measured with a pH probe (n = 4). The dashed purple and yellow lines show the upper and lower detection limits of the pH dye, respectively.

When growing alone (Fig. S12), *C. freundii* showed similar growth at all antibiotic concentrations, and acidified more but grew worse at increasing glucose concentration, suggesting that it also suffered from its own acidification. *P. aeruginosa* instead grew worse at higher ampicillin concentrations, but was able to recover and survive until the end of the experiment in almost every glucose and ampicillin concentration when alone (Fig. 3A, top). When co-cultured with *C. freundii*, however, *P. aeruginosa* dropped below the detection limit in all replicate populations at ampicillin concentrations of 800*µ*g/mL and in all replicates at glucose concentrations of 30mM. *P. aeruginosa* survival was highly dependent on pH, as *P. aeruginosa* was able to grow whenever the pH remained above 6, as long as the ampicillin concentration was low enough (Fig. 3B). Population recovery occurred only once at 26mM of glucose and not a single time at 30mM of glucose, where *C. freundii* was able to acidify the environment consistently, even when no ampicillin was added. In sum, from the model we had expected a higher population density of *C. freundii* or a higher glucose concentration to lead to faster acidification, acting synergistically with the antibiotics to inhibit *P. aeruginosa*. Our transfer experiments showed that – as long as we pipetted the bacterial culture often enough – *P. aeruginosa* could be eliminated after the first transfer if the pH dropped sufficiently. The drop in pH was possible if the glucose concentration crossed a threshold beyond which the pH could drop below 6. When grown alone, the pH remained high such that *P. aeruginosa* survived at all glucose concentrations, and could only sometimes be eliminated with high concentrations of ampicillin.

### Model predicts that this approach can be applied to other species pairs

Having shown that acidification by *C. freundii* at high glucose concentrations could effectively eliminate *P. aeruginosa*, we wondered whether this could work for other species pairs. To evaluate the generality of our approach, we first used the mathematical model to guide our expectations. This time, in addition to varying the initial antibiotic concentrations, we explored how different properties of the species would change the outcome. For example, in the original model we parameterized the pathogen to grow slightly faster than the commensal (0.45 versus 0.4) to approximate our experimental data. The commensal was also more resistant to the antibiotic than the pathogen and had a higher tolerance for acidity. Would our “feed your enemy’s enemy” approach still work if our species pairs did not meet these criteria? And which criteria are needed for success?

The model predicts that the most important properties of the pathogen and commensal species are that: (i) the commensal does not grow much slower than the pathogen on the provided carbon source (Fig. 4A) and (ii) the commensal needs to have a higher tolerance to acidity compared to the pathogen (Fig. 4D, E). If either of these criteria are not met (i.e. if the pathogen grows a lot faster than the commensal or the commensal is not much more tolerant to low pH), more antibiotic will be needed to eliminate the pathogen. Whether or not the pathogen consumes the organic acid and the resistance of the commensal to the antibiotic were less important according to our model (Fig. 4B, C). This last point was encouraging, as it suggests that one need not use a resistant commensal strain, in which case the antibiotic can be completely omitted. Indeed, applying the antibiotic can decrease the efficacy of the approach in these cases.

**Fig. 4.**
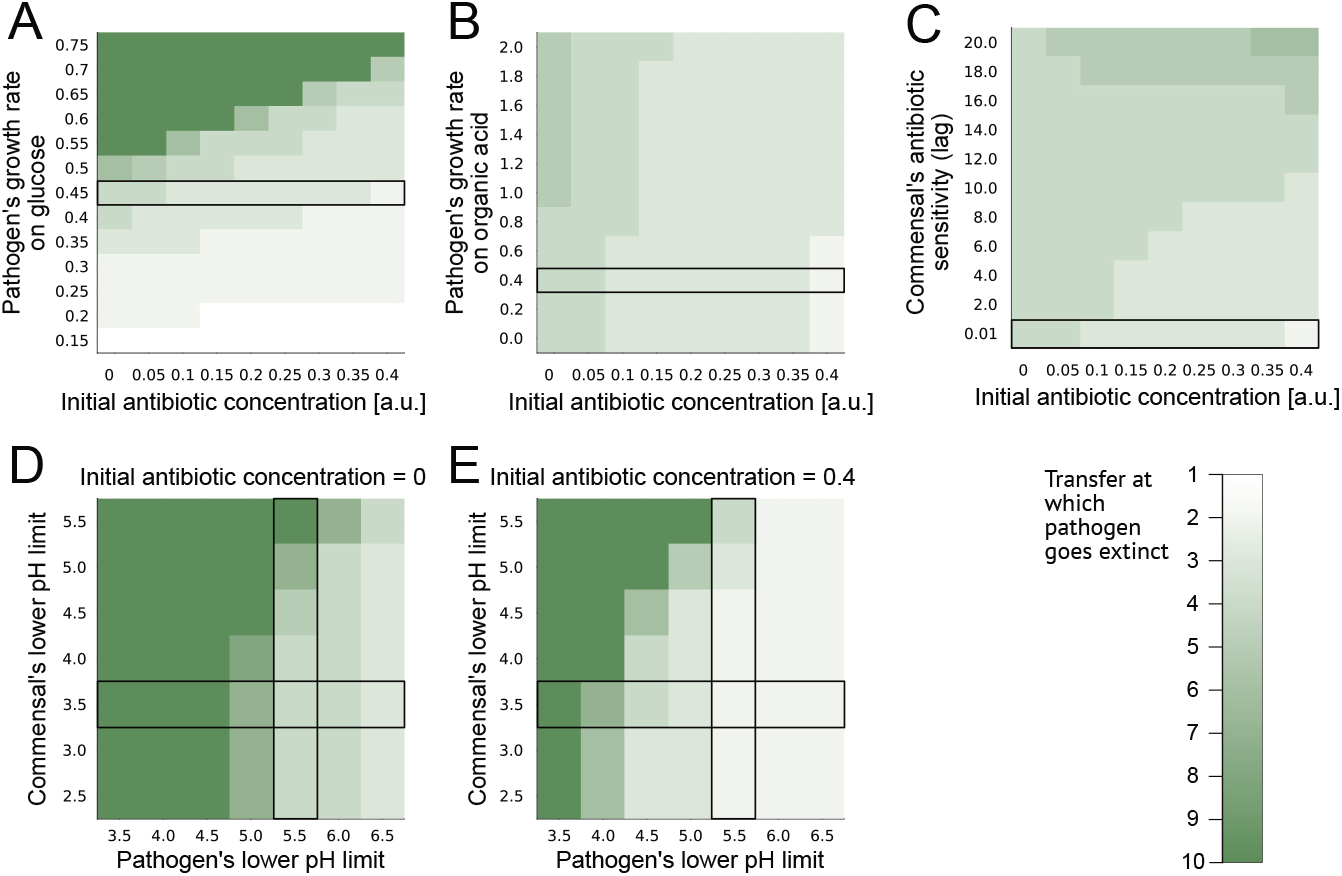
Testing “feed your enemy’s enemy” across a range of species parameters. Predictions generated by the mathematical model varying different properties of the commensal and pathogen. Black rectangles in each plot show the default parameters used in the simulations shown in Fig. 2. The shade of green shows the transfer at which the pathogen goes extinct, with 10 meaning it doesn’t until the end of the simulations. (A) If the pathogen grows significantly faster than the commensal (0.45 in these simulations) on the main supplied carbon source (glucose), more antibiotic will be needed to eliminate it. (B) Its growth rate on the resulting organic acid does not affect the results much. (C) It helps if the commensal is more resistant to the antibiotic, but even if it is not, the pathogen can be eliminated. If too much antibiotic is added and the commensal is sensitive, the approach works less well. (D) The pathogen needs to be less tolerant to the pH compared to the commensal in the absence of antibiotics. (E) Adding antibiotics (concentration 0.4) can compensate for having more similar tolerance levels.

### Testing other species pairs experimentally

We next tested these predictions experimentally by exchanging *P. aeruginosa* for two different pathogens, *Agrobacterium tumefaciens*, a plant pathogen and *Klebsiella pneumoniae*, another opportunistic human pathogen. We also tested a different commensal: *Lactobacillus plantarum*, a commonly used probiotic species.

*K. pneumoniae* had a negative effect on the growth of *C. freundii* in the absence of ampicillin. But because of the higher resistance of *C. freundii* to ampicillin, when 800 *µ*g/mL of ampicillin were added, *K. pneumoniae* grew significantly worse in co-culture compared to mono-culture (Fig. 5A bottom). Strong acidification was observed in the mono-cultures of *C. freundii* and in the co-culture with ampicillin. This suggests that *K. pneumoniae* would be eliminated more easily in transfers with ampicillin and *C. freundii* than with ampicillin alone due to the acidification of the medium. Ampicillin would nevertheless be needed in this case.

**Fig. 5.**
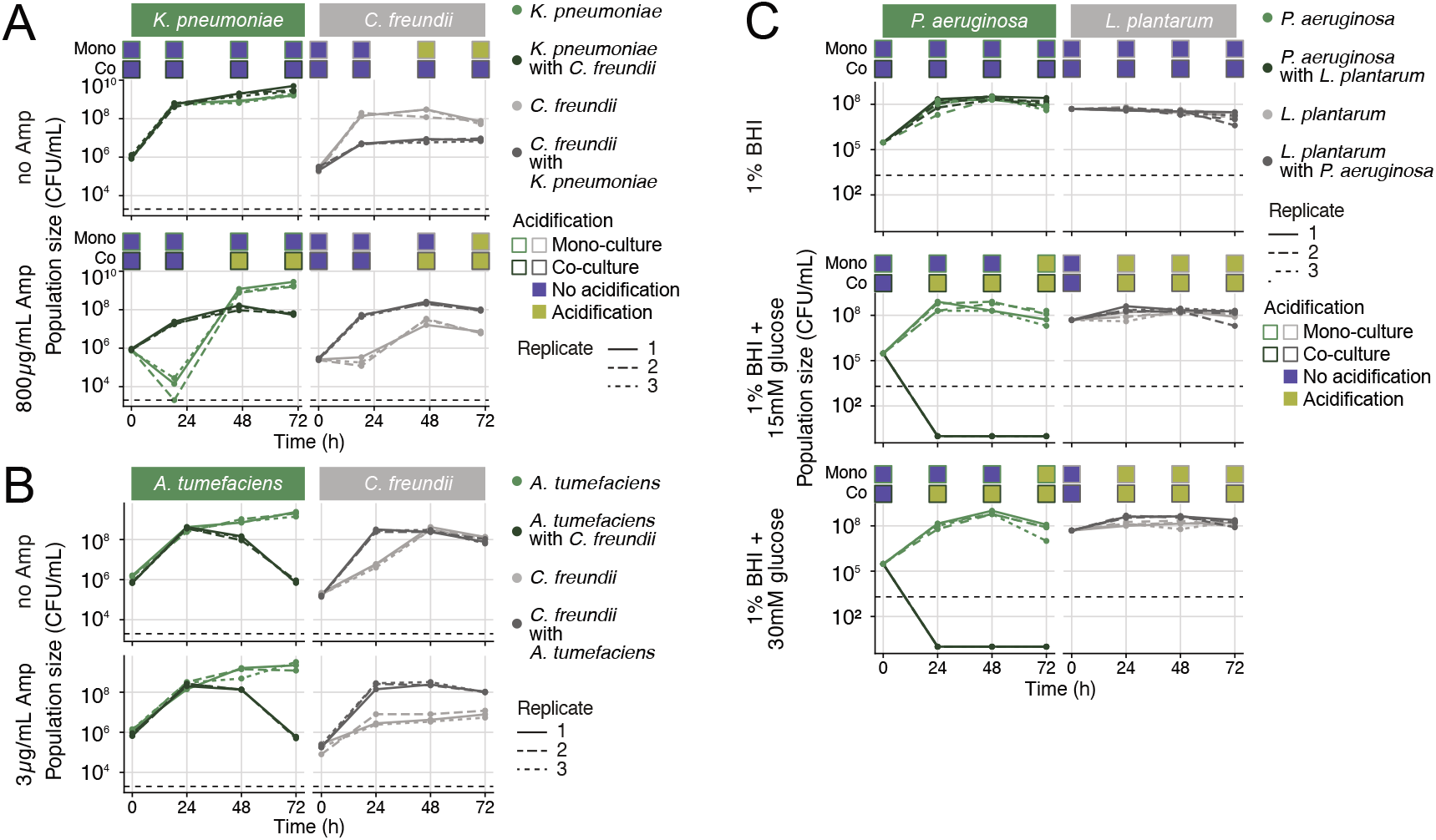
Testing “feed your enemy’s enemy” with other species pairs experimentally. (A) *C. freundii* (selected for increased resistance to ampicillin) can inhibit the growth of *K. pneumoniae* in co-culture at 800*µ*g/mL of ampicillin. Without ampicillin, it is *C. freundii* that grows worse. (B) *C. freundii* (WT, i.e. not selected for increased resistance to ampicillin) can inhibit *A. tumefaciens* regardless of the presence of ampicillin. (C) *P. aeruginosa* is inhibited by *Lactobacillus plantarum* but only if glucose is added to BHI medium. Even without antibiotics, *P. aeruginosa* does not survive until the end of what would be the first transfer (as opposed to with *C. freundii* in Fig. 1C, Fig. 3). pH measurement (when applicable) is shown in purple and yellow boxes above each measured time-point.

*A. tumefaciens* was also inhibited by *C. freundii* compared to when it was grown alone. In this case, the effect was independent of the presence of ampicillin. As this experiment was one of the first ones we conducted, the original wildtype *C. freundii* was used, which had an MIC of 3 *µ*g/mL, and we did not measure the pH. Nevertheless, this experiment suggested that *A. tumefaciens* would be eliminated in a co-culture with *C. freundii*, which we confirmed in transfer experiments (Fig. S13), where *A. tumefaciens* was eliminated faster in the absence of ampicillin (in transfer 2 versus transfer 4 at 3 *µ*g/mL of ampicillin), as predicted by the model for a more sensitive commensal (Fig. 4C).

Next, we changed the commensal to *Lactobacillus plantarum*, a common probiotic species that is a known to acidify by producing lactic acid (24). We did not manage to increase the resistance of *L. plantarum* to antibiotics, nor to grow it in the minimal medium used above. Given our model predictions, though, we anticipated that it would be worth testing its effect on the pathogens even without antibiotics in rich medium with increasing glucose concentrations. As of 15mM of glucose, *L. plantarum* inhibited *P. aeruginosa* completely, such that no cells were detected after 24h of a batch co-culture (Fig. 5C). This effect is striking, given that alone, *P. aeruginosa* could grow to an approximate population size of 10^8^ CFU/ml. In co-culture with *K. pneumoniae* the effects were more variable (Fig. S14): The first time we ran the experiment, *K. pneumoniae* dropped below the detection limit by 48 or 72 hours, at glucose concentrations 30mM or higher. But when we repeated the experiments two more times, *K. pneumoniae* was not or barely affected by *L. plantarum*. This suggests that antibiotics may be needed in addition to the acidifying species to more robustly suppress *K. pneumoniae* (as in Fig. 5A).

Overall, our model and experiments demonstrate that although different species will have different properties such that the precise conditions in which our approach works will differ, it is in principle not restricted to *C. freundii* and *P. aeruginosa*. We do expect though that in some cases, for example, if the pathogen is highly competitive or the commensal too sensitive to low pH, it will not be possible to eliminate the pathogen.

## Discussion

How the chemical environment shapes interactions between bacterial species is critical not only to understand microbial community dynamics and function generally, but also to control a species of interest. In previous work, we had shown how to switch the interactions between two bacterial species from negative to positive by changing the chemical environment (20), while in this study we apply the control of interactions to eliminate a pathogen through the increase of glucose concentration together with – or instead of – antibiotic administration. This works by feeding the acidifier, i.e. “feed the enemy’s enemy”, to reduce the pH thereby inhibiting pathogens that are less sensitive to low pH. Importantly, by testing this principle on several commensal-pathogen pairs, we demonstrate the generality of our idea.

These observations build on and connect to the idea that acidification promotes pathogen colonization resistance, which is important in the context of the host microbiome (10) and in food preservation (19). The idea that the drop in pH can be mediated by resource concentrations to strengthen colonization resistance has, instead, only rarely been shown (25).

Our work also relates to studies that have focused on how interactions between species, such as cross-feeding (9, 26) or cross-protection (7, 27) can reduce the efficacy of antibiotic treatment (28–30), except that in this case, we are exploiting negative interactions between species. We also know that antibiotics and low pH (abiotically) can have synergistic effects in several bacterial species and contexts (21–23, 31–33), as pH is often modified by bacteria and is of great importance to homeostasis (16, 34) and can change the conformation of transport proteins of bacterial membranes (17, 18).

In an applied medical context, the ultimate goal is to reduce the concentrations of antibiotics used to eliminate pathogens. We consider our approach to be highly relevant to counter the rise of antibiotic resistance. In practice, we are proposing to administer, in addition to antibiotics, an acidifying probiotic together with additional resources (prebiotics, e.g. glucose) that would increase acidification. Alternatively, one could evaluate the presence of pre-existing pH-modulating species in the host and stimulate their growth by administering prebiotics. This approach would be particularly interesting for combating plant pathogens like *A. tumefaciens* (Fig. 5B, S13) where the use of antibiotics is restricted in many countries. Applying “feed the enemy’s enemy” in the context of a host is still a few steps away, though. In the host, pH changes might be tightly buffered and controlled (35), meaning that a commensal may never be able to acidify the environment significantly. Another complexity of a real environment is that other microbial species are likely to also be present and may be affected positively or negatively by the changes in pH and in turn also affect the targeted pathogen. Nevertheless, in the spatially structured environment of the host tissue, it has been suggested that acidification can act locally to inhibit pathogens (11).

Two additional factors can be important in shaping interactions and would be interesting to explore in future work: the role of spatial organization, and the evolution of resistance (Fig. S7). Regarding spatial structure, the extinction of *P. aeruginosa* depended on frequently mixing and physically disturbing the cultures, while leaving them undisturbed almost eliminated the antibiotic effect (Fig. S4, S5). Our current hypothesis is that *P. aeruginosa* may have formed biofilms in the undisturbed cultures as a response to antibiotic stress (36), which protected them against the antibiotic (37, 38). *P. aeruginosa* also forms more biofilm and gains resistant faster in the acidic conditions of cystic fibrosis lungs (39), which corroborates our observations.

Our findings highlight the importance of considering the community and chemical context of pathogens during antibiotic treatment of microbial infections. In most cases, the outcome of antibiotic treatment is driven by several dynamic variables that are the direct result of bacterial interactions and growth, including the pH. We demonstrate that the strength of these interactions, such as the degree of acidification, can be externally controlled by manipulating the chemical environment, in this case the addition of glucose. A deeper understanding of such variables is key to controlling microbial interactions, thereby providing an alternative strategy to enhance or in some cases even replace the use of antibiotics. Our work advocates for studying the effect of antimicrobial compounds in environmental conditions that better mirror the complexity of bacterial communities (40), particularly in this broader framework of community control so that we can increase our ability to obtain successful treatment outcomes.

## Material and methods

### Bacterial strains

The following bacterial species were used in this study: *Pseudomonas aeruginosa* PA14 (kindly donated by Kevin Foster), *Citrobacter freundii* and *Klebsiella pneumoniae* (kindly donated by Peter Küenzi), *Agrobacterium tumefaciens* (kindly donated by Christopher van der Gast and Ian Thompson), and *Lactobacillus plantarum* (kindly donated by Julian Garneau and Pascale Vonaesch). We used a strain of PA14 that contains an mCherry fluorescent tag. We used two strains of *C. freundii*: the strain used in the experiments together with *P. aeruginosa* and *K. pneumoniae* was selected for increased resistance on ampicillin medium. The original isolate that was more sensitive to ampicillin is used in experiments with *A. tumefaciens* (designated as “WT”).

### Cell cultures

To start bacterial cultures, a single colony of the strain of interest was grown overnight (200 rpm, 28°C) in 10 mL of TSB (tryptic soy broth) and then sub-cultured for 3 hours in the same medium at an initial OD_600_ of 0.05. Unless mentioned otherwise, the OD_600_ was adjusted to 0.1 and then the culture was washed 3 times with PBS (phosphate buffer saline) and then resuspended in minimal medium (concentration of glucose depends on the experiment). Finally, this bacterial culture was used to inoculate the different conditions of our experiments, diluting it 100× (final OD_600_ = 0.001). For co-cultures, a final OD_600_ of 0.001 of each species were combined, such that mono-and co-cultures contained similar population sizes of each species. When *C. freundii* was 10× more, the starting culture was OD_600_ of 1 (final OD_600_ = 0.01). For *L. plantarum*, cells were pre-cultured in Brain Heart Infusion (BHI) instead of TSB. *L. plantarum* cultures were always set to an initial OD_600_ of 0.001, as was *P. aeruginosa* when co-cultured with *L. plantarum*.

Growth and dilution cycles were performed by diluting bacterial cultures every 72h 100× into fresh medium corresponding to the same initial glucose and ampicillin concentrations.

### Media recipes

For all cultures involving species other than *Lactobacillus plantarum*, we used an M9 minimal medium with glucose. To make 1L of medium, we mixed 100mL of 10 × M9 (pH adjusted to 7.4 with NaOH), 20mL of 50 × HMB (see (20) for recipes), glucose (e.g. 40mL of a 0.75M stock solution to achieve a final concentration of 30mM) and completed the volume with water. TSB and BHI were purchased from BD Difco.

### Measuring population sizes

To measure population sizes, we quantified CFUs. We first sampled 20 *µ*g/mL of bacterial culture and serially diluted it down to 10^−7^ (in V = 200 *µ*l). A volume of 5 *µ*l was plated in the form of a drop on TSA (tryptic soy agar) for each 10-fold dilution. Plates were incubated at 28°C. For cocultures, TSA allowed to distinguish *P. aeruginosa* and *C. freundii* as i) the strains have distinct colors (*P. aeruginosa* is pink and *C. freundii* is white) as *P. aeruginosa* has an mCherry tag and ii) *C. freundii* colonies grow after 12 hours while *P. aeruginosa* takes between 24 and 48 hours to show countable colonies. *L. plantarum* grew slowly on TSA, and its colonies were instead counted on MRS plates. The limit of detection is at 10^2^ CFUs/mL.

We also quantified bacterial densities in some cases by measuring optical density (OD) at 600 nm in a micro-plate reader (BioTek Synergy H1 at 28°C with continuous double-orbital shaking), every 30 min for 72 hours (continuous shaking, 28°C).

### MIC and growth parameter estimation

In a 96-well plate, the first well of each line was filled with 200 *µ*L of an ampicillin stock (Ampicillin Sodium Salt *BioChemica*, code A0839, ampicillin powder was diluted in minimal medium with glucose, either 15 mM or 30 mM depending on the experiment). The rest of the wells were filled with minimal medium with glucose (at equal concentrations of glucose, see below). The ampicillin was serially diluted 2x until the penultimate well (final volume = 100 *µ*g/mL). The bacterial culture was prepared as described above, but at a final OD_600_ of 0.2 in minimal medium with glucose, which was then diluted 100 times in the same medium to obtain a stock culture. We added 100 *µ*L of this stock to all wells of the 96-well plate so that all wells contained bacteria at an initial OD_600_ of 0.001 and either 15 mM or 30 mM of glucose (V = 200 *µ*g/mL). Growth was followed by measuring OD_600_ over time as described above. Lag length and final yield (Fig. S1) were determined manually: final yield is OD_600_ at 72h and lag length is the timepoint at which the OD_600_ passed a threshold value of 0.09.

### MICs over a range of pH values

In a 96-well plate, the first well of each line was filled with 200 *µ*L of ampicillin stock (800 *µ*g/mL in water). The rest of the wells were filled with 100 *µ*L of water. We then proceeded to a serial 2x dilution until the penultimate well (final volume = 100 *µ*g/mL). The bacterial cultures were prepared as described above, however we prepared 6 bacterial stock cultures at OD = 0.002: one per pH condition tested. We thus prepared minimal media with 15 mM of glucose and various concentrations of HCl to obtain a range of pH levels (see table 1, media concentrated 2x to account for the 2x dilution in the 96-well plate). Each stock culture was then added to one dilution line of the 96-well plate (100 *µ*L per well). Growth was followed by measuring OD_600_ over time as described above.

**Table 1.**
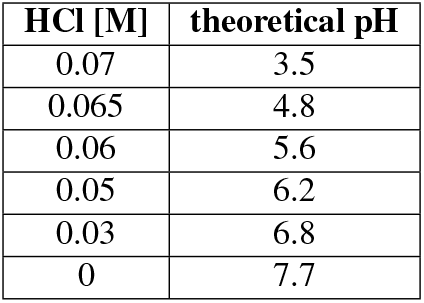
Concentration of HCl added to minimal medium with glucose to obtain a range of pH levels from neutral to acidic.

| HCl [M] | theoretical pH |
| --- | --- |
| 0.07 | 3.5 |
| 0.065 | 4.8 |
| 0.06 | 5.6 |
| 0.05 | 6.2 |
| 0.03 | 6.8 |
| 0 | 7.7 |

### Use of bromocresol purple to determine the pH of cell cultures

To follow the pH of bacterial cultures over time without using a pH-meter, we used a pH dye: bromocresol purple. This dye is purple at pH ≥ 6.8 and yellow at pH ≤ 5.2. The purple form of the dye absorbs at 588 nm and can easily be correlated to the pH value of the medium it is in. The absorption of the dye being so close to 600 nm (absorption of bacterial cells), we measured the density of bacterial cells at 750 nm. We did an absorption scan on *P. aeruginosa* and *C. freundii* which showed that they absorb similarly at 600 and 750 nm. Therefore, we can calculate the net absorbance of the bromocresol purple by measuring the absorbance at 588 and 750 nm and then subtracting the OD_750_ from the OD_588_. From that value, we can calculate the pH by establishing a standard curve to relate the intensity of the absorption at 588 nm to the pH. To do so, we prepared several citrate-Na_2_HPO_4_ buffer solutions (see table 2) to have a range of solutions with known stable pH levels (from pH 4.8 to 7.2). In a 96-well plate, we put 180 *µ*L of each buffer and added 20 *µ*L of bromocresol pruple solution 0.01% (w/w) (in triplicate). We measured the OD_588_, performed a linear regression and extracted the equation to calculate the pH from the absorbance.

**Table 2.**
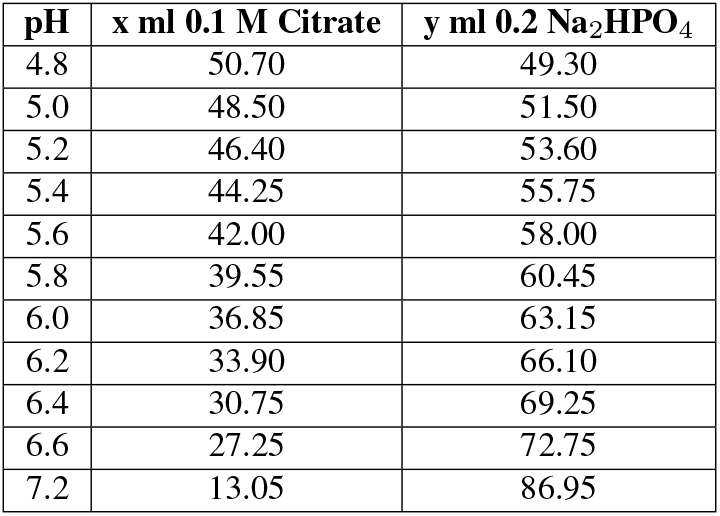
Citrate-Na_2_ HPO_4_ buffers in a range of pH levels from neutral to acidic used to produce the standard curve of bromocresol purple at 588 nm.

### Mathematical model

The mathematical model is a consumer-resource model based on ordinary differential equations, describing changes in the abundances of the two species, concentrations of glucose and organic acid, and the pH. It is described in detail in the supplement (note S1). The code is available on github here.

## Data and code availability

All experimental data and code are available on Zenodo (DOI: 10.5281/zenodo.17751754).

## Acknowledgments

We thank Julien Luneau, Eric Ulrich, Prajwal Padmanabha and Margaret Vogel for useful and constructive feedback on the manuscript. We thank Kevin Foster, Julian Garneau, Pascale Vonaesch, Christopher van der Gast, Ian Thompson and Peter Küenzi for providing the strains used in this study.

