## Supplementary methods and figures for "Feeding your enemy’s enemy: Acidifying bacteria inhibit pathogenic bacteria more strongly with increasing glucose"

### Supplementary Note S1: Mathematical model

To better understand the dynamics that we observed within our 2-species community (Fig. 1), we built a consumer-resource model based on modified Monod equations (with maximum growth rate $r$, half-saturation constant $K$ and yield $Y$). In our model, species $B_1$ (pathogen) and $B_2$ (commensal) can grow together using the resource $C$ and produce a certain quantity of organic acid $A$ that is proportional to the quantity of $C$ consumed. Species $B_1$ produces a very small quantity of $A$ while $B_2$ produces a significant quantity of $A$ (described by the parameter $a_i$) that will acidify the medium through the net production of protons $H$ (directly used to calculate the pH using equation S6). Species $B_1$ can consume a fraction $q$ of organic acid $A$ through cross-feeding. The value of $H$ is obtained after applying a neutralisation function (equation S7) that mimics the effect of a chemical buffer (used in our growth medium) on the concentration of organic acid $A$. Species $B_1$ and $B_2$ have a sensitivity to acidic pH that is described by equation S8 (41) that depends on 2 parameters: $pK_h$ and $pK_l$ which are respectively the high and low pH bounds, i.e. the pH value above and below which the species will start being sensitive to the pH, respectively. By default, species $B_1$ is sensitive to pH below 5.5 and $B_2$ below 3.5. In addition, there is a concentration $b$ of antibiotic that affects $B_1$ and $B_2$ and depends on the sensitivity parameter $l_i$, which induces a lag in bacterial growth. $B_1$ has a high sensitivity to ampicillin while $B_2$ is mostly resistant.

#### System of ordinary differential equations (ODEs).

$$\frac{dB_1}{dt} = r_{1,1}\frac{C}{C+k_{1,1}}\frac{t^2}{(l_1b)^2+t^2}F_1(pH(H))B_1 + r_{1,2}\frac{qA}{qA+k_{1,2}}\frac{t^2}{(l_1b)^2+t^2}F_1(pH(H))B_1 \quad \textbf{(S1)}$$

$$\frac{dB_2}{dt} = r_{2,1}\frac{C}{C+k_{2,1}}\frac{t^2}{(l_2b)^2+t^2}F_2(pH(H))B_2 \quad \textbf{(S2)}$$

$$\frac{dC}{dt} = -\frac{1}{Y_{1,1}}\frac{r_1}{C+k_1}\frac{t^2}{(l_1b)^2+t^2}F_1(pH(H))CB_1 - \frac{1}{Y_{2,1}}\frac{r_2}{C+k_2}\frac{t^2}{(l_2b)^2+t^2}F_2(pH(H))CB_2 \quad \textbf{(S3)}$$

$$\frac{dA}{dt} = (\sum a_i\frac{1}{Y_i}\frac{r_i}{C+k_i}\frac{t^2}{(l_ib)^2+t^2}F_i(pH(H))CB_i) - \frac{1}{Y_{1,2}}\frac{r_{1,2}}{qA+k_{1,2}}\frac{t^2}{(l_1b)^2+t^2}F_1(pH(H))qAB_1 \quad \textbf{(S4)}$$

$$\frac{dH}{dt} = N(pH(H))(1-q)\frac{dA}{dt} \quad \textbf{(S5)}$$

Where the pH is:

$$pH(H) = \frac{1}{2}(pKa - log(H)), \quad \textbf{(S6)}$$

which is the general formula to calculate the pH of a weak acid. Here the $pKa$ always takes the value 9.

The neutralisation (buffering) function is:

$$N(pH(H)) = \frac{1}{1+e^{-m(pH(H)-x_0)}} \quad \textbf{(S7)}$$

This logistic function depends on the pH and takes a value between 0 and 1 depending on the pH (0 = maximum buffering capacity, 1 = no buffering capacity). The change in organic acid $A$ is multiplied by this term, which cancels a fraction of $A$, thereby buffering the medium. Here, $m$ always takes the value $-5$ and $x_0$ always takes the value 5.5.

The pH inhibition function is defined as (from (41)):

$$F_i(pH(H)) = \frac{1+2*10^{0.5(pK_{li}-pK_{hi})}}{1+10^{(pH-pK_{hi})}+10^{(pK_{li}-pH)}} \quad \textbf{(S8)}$$

This bell-shaped function depends on the pH and can take values between 0 and 1 (0 = full inhibition, 1 = no inhibtion). The growth of $B_1$ or $B_2$ is multiplied by this function, which cancels a fraction of growth.
The model was implemented in Julia 1.8.5. ODEs were solved with the package `DifferentialEquations` using the method `Rosenbrock23`, which is an order 2/3 L-Stable Rosenbrock-W method, appropriate for very stiff equations with oscillations at low tolerances.

| Default species-specific parameters | | $B_1$ | | |
|---|---|---|---|---|
| **Parameter** | **Description** | $B_1$ **(i = 1) - Glucose (j = 1)** | $B_1$ **(i = 1) - Organic acid (j = 2)** | $B_2$ **( i = 2)** |
| $r_{i,j}$ | growth rate | 0.45 | 0.45 | 0.4 |
| $k_{i,j}$ | half saturation constant | 8 | 5 | 8 |
| $Y_{i,j}$ | yield | 10000 | 40000 | 10000 |
| $l_i$ | lag factor | 15 | | 0.01 |
| $a_i$ | acidification rate | 10^(-15) | / | 0.2 |
| $pK_{li}$ | lower pH bound | 5.5 | | 3.5 |
| $pK_{hi}$ | higher pH bound | 8.5 | | 8.5 |

**Table S1.** List of all species specific parameters and their values

| Default global parameters | | |
|---|---|---|
| **Parameter** | **Description** | **Value** |
| $b$ | antibiotic concentration | 0.3 |
| $q$ | fraction of cross-fed organic acid | 0.1 |
| $t$ | time | from simulation |
| $pKa$ | acid dissociation equilibrium | 9 |
| m | slope of neutralization | -5 |
| $x_0$ | midpoint of neutralization | 5.5 |

**Table S2.** List of all global parameters and their values.

### Supplementary figures

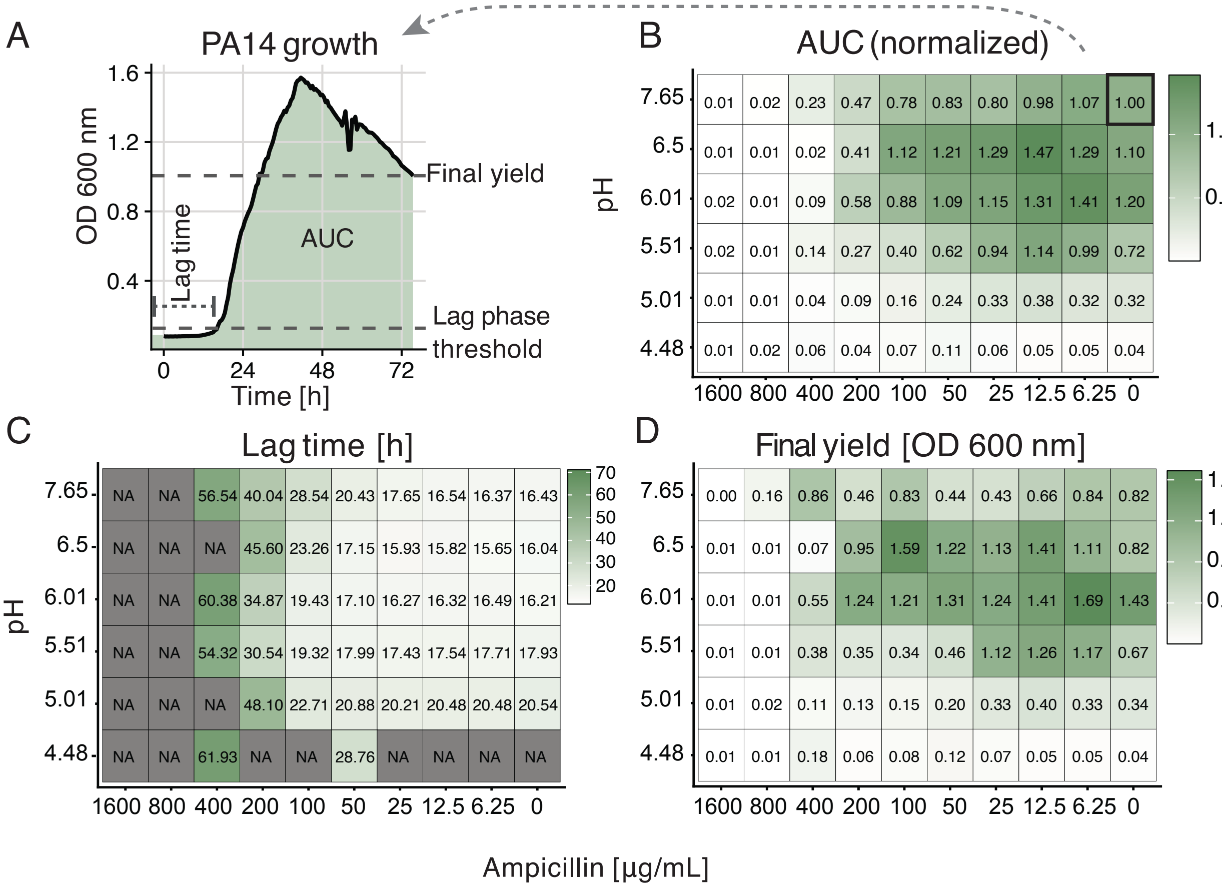


**Fig. S1.** Effect of ampicillin combined to acidification on *P. aeruginosa*. (A) Growth of *P. aeruginosa* in non-acidified medium without ampicillin. The different growth parameters considered in the following panels are displayed. An ampicillin- combined with pH-sensitivity assay was performed and the normalized area under the growth curve (AUC) (B), the lag time (C) and the final yield (D) were determined. “NA” in panel (C) indicates “not applicable”, i.e. the $OD_{600}$ never exceeded the threshold, here: 0.09.

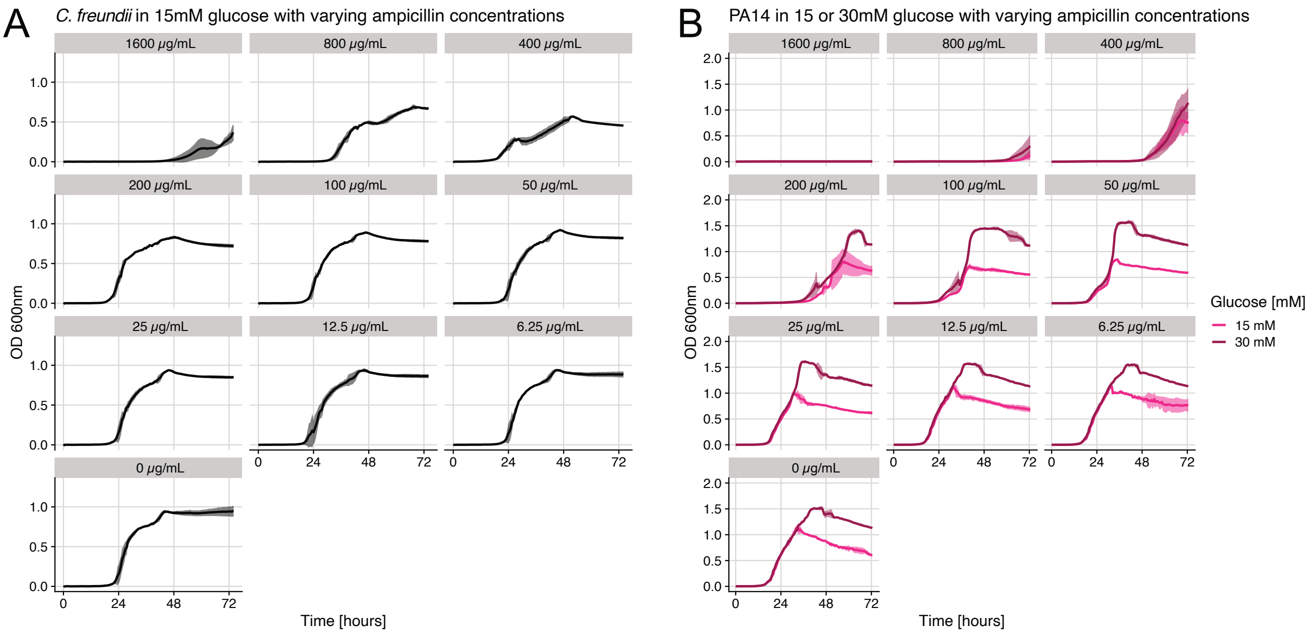


**Fig. S2.** (A) Growth of *C. freundii* (isolate selected for resistance against ampicillin, not WT) in varying concentrations of ampicillin in minimal medium with 15mM of glucose. (B) MIC of *P. aeruginosa* in varying concentrations of ampicillin in minimal medium with either 15 or 30mM of glucose. For all curves, $OD_{600}$ was measured every 30 min for approximately 72 hours (n = 3).

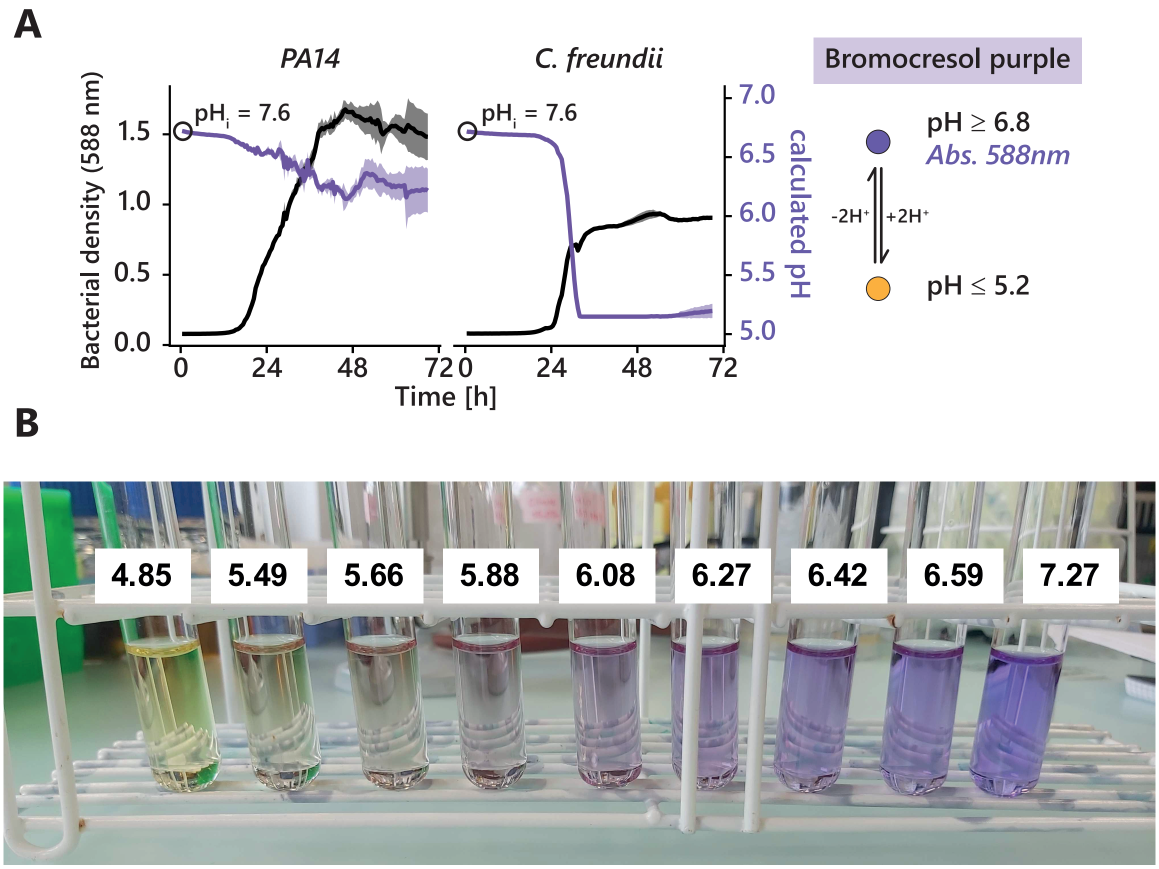


**Fig. S3.** Use of the pH dye bromocresol purple to determine the pH of bacterial cultures. (A) Growth curve and pH curves of *P. aeruginosa* and *C. freundii* grown in minimal medium with 30 mM of glucose. The medium contained 0.001 % (w/w) of bromocresol purple (BP). Bacterial density was measured at 750 nm and the absorbance of BP was measured at 588 nm and corrected to remove bacterial density, (n = 3). For more details see methods. (B) Color range of BP in cell-free liquid with pH adjusted as indicated.

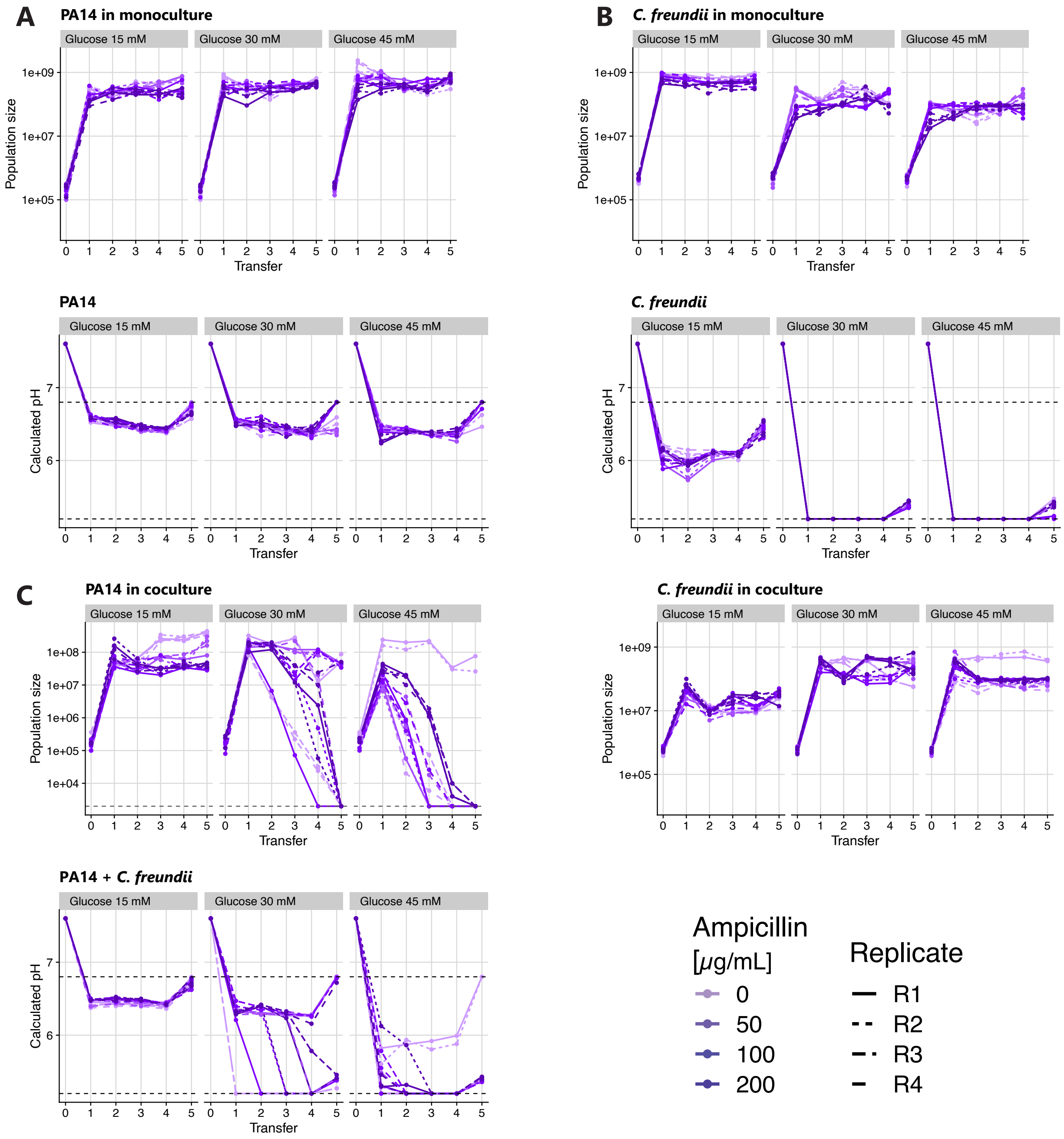


**Fig. S4.** Approximately equal initial population size of *P. aeruginosa* and *C. freundii* (both initial $OD_{600}$ of 0.001), antibiotic gradient up to 200$\mu$g/mL. Growth of *P. aeruginosa* (A) and *C. freundii* (B) over 5 transfers (3 days of growth, 1 % transfer in fresh medium, CFUs at each transfer) in mono-culture (top row) and pH value at each timepoint from the absorbance of bromocresol purple measured at $OD_{588}$ at each transfer (bottom row). (C) Growth of *C. freundii* and *P. aeruginosa* when they are co-cultured as well as the pH values (same as in Fig. 3C).

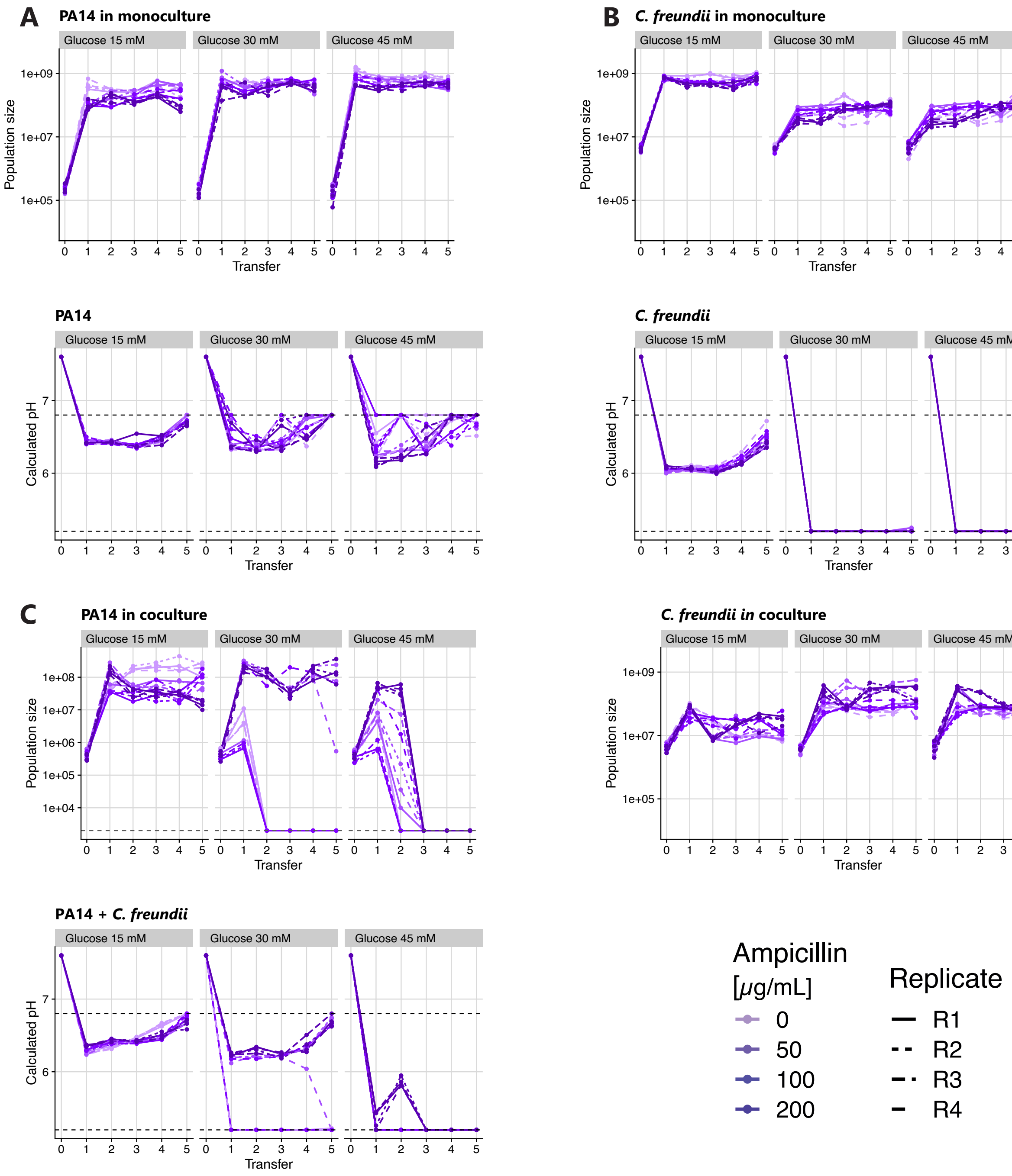


**Fig. S5.** Same as the experiment shown in Fig. S4 but the initial $OD_{600}$ of *C. freundii* (0.01) is 10× higher than *P. aeruginosa* (0.001), antibiotic gradient up to 200$\mu$g/mL. Growth of *P. aeruginosa* (A) and *C. freundii* (B) over 5 transfers (3 days of growth, 1 % transfer in fresh medium, CFUs at each transfer) in mococulture (upper graph) and pH value at each timepoint from the absorbance of bromocresol purple measured at $OD_{588}$ at each transfer (lower graph). (C) Growth of *C. freundii* and *P. aeruginosa* when they are cocultured as well as the pH values (same as in Fig. 3C).

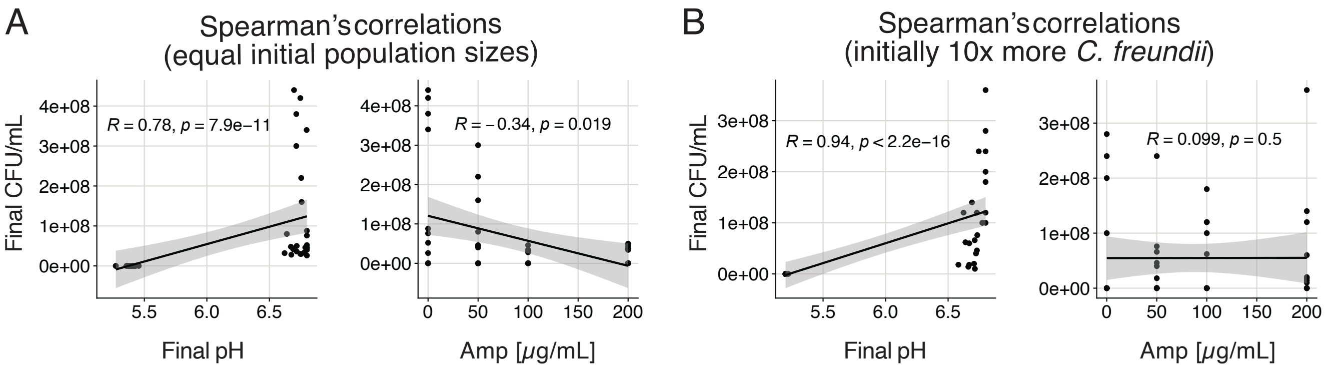


**Fig. S6.** **(A)** From the data in Fig. S4, Spearman correlations between the final population size and the final pH (transfer 5) (left) and the ampicillin concentration (right). **(B)** From the data in Fig. S5, Spearman correlations between the final population size and the final pH (transfer 5) (left) and the ampicillin concentration (right).

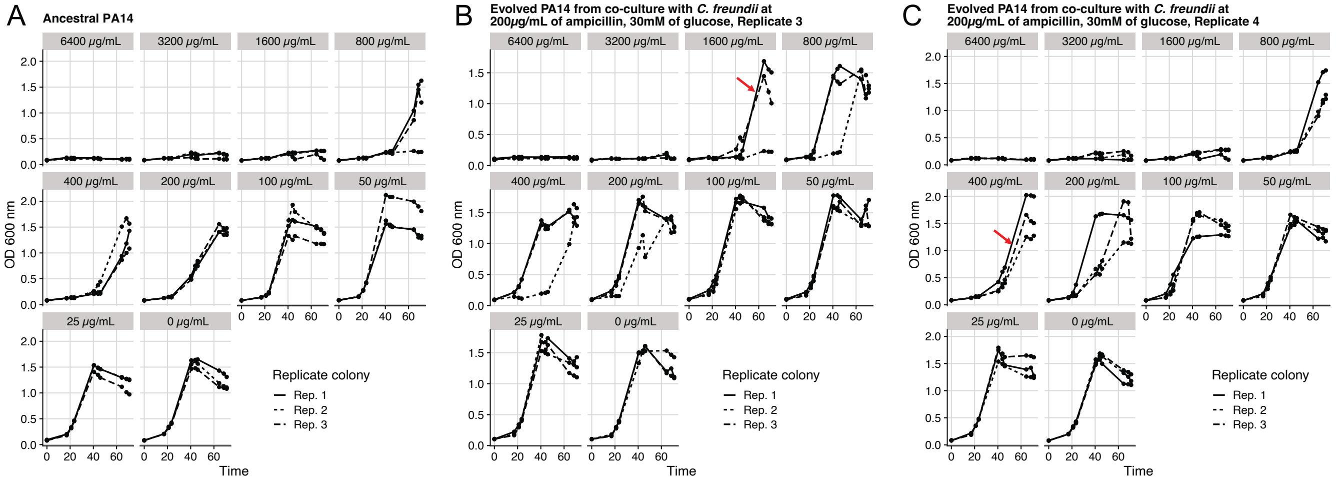


**Fig. S7.** MIC against ampicillin of (A) the ancestral strain of *P. aeruginosa* compared to some of the isolates from the fifth transfer in two experimental replicates (B) and (C) where they had survived in co-cultured with *C. freundii* at 30mM of glucose and 200$\mu$g/mL of ampicillin. We show three individual colonies that were isolated and grown separately in minimal medium with 30 mM of glucose and the $OD_{600}$ was measured 3 times per day for 3 days in a micro-plate reader (n = 3). While the ancestral *P. aeruginosa* has an MIC of 800$\mu$g/mL, some of the evolved isolates, but not all, showed a higher MIC of up to 1600$\mu$g/mL. Increased resistance to ampicillin is highlighted in the curves indicated with red arrows. Resistance was not universal, though, showing that survival could also be due to tolerance.

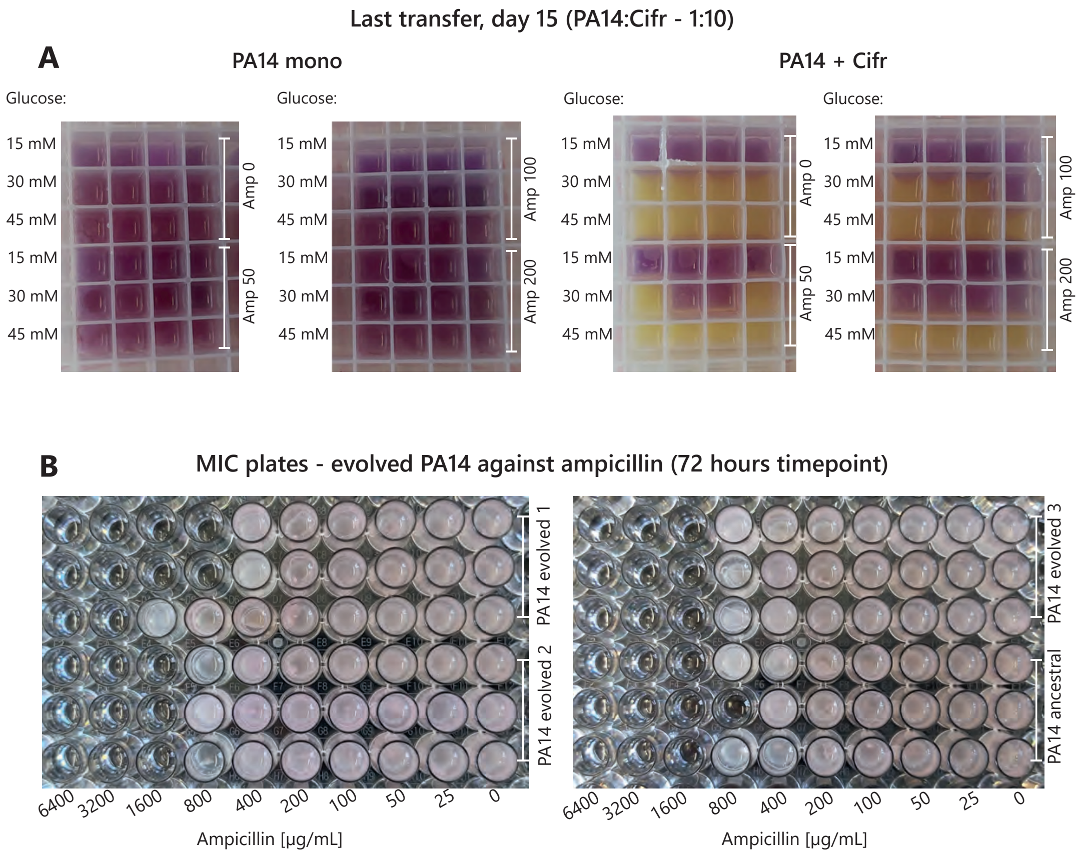


**Fig. S8.** Examples of biofilms in *P. aeruginosa* cultures. (A) Pictures of the last timepoint of the transfer experiment in Fig S5A, C. On the left is *P. aeruginosa* in monoculture, and on the right in coculture with *C. freundii*. (B) Pictures of the MIC assay plates from Fig. S7.

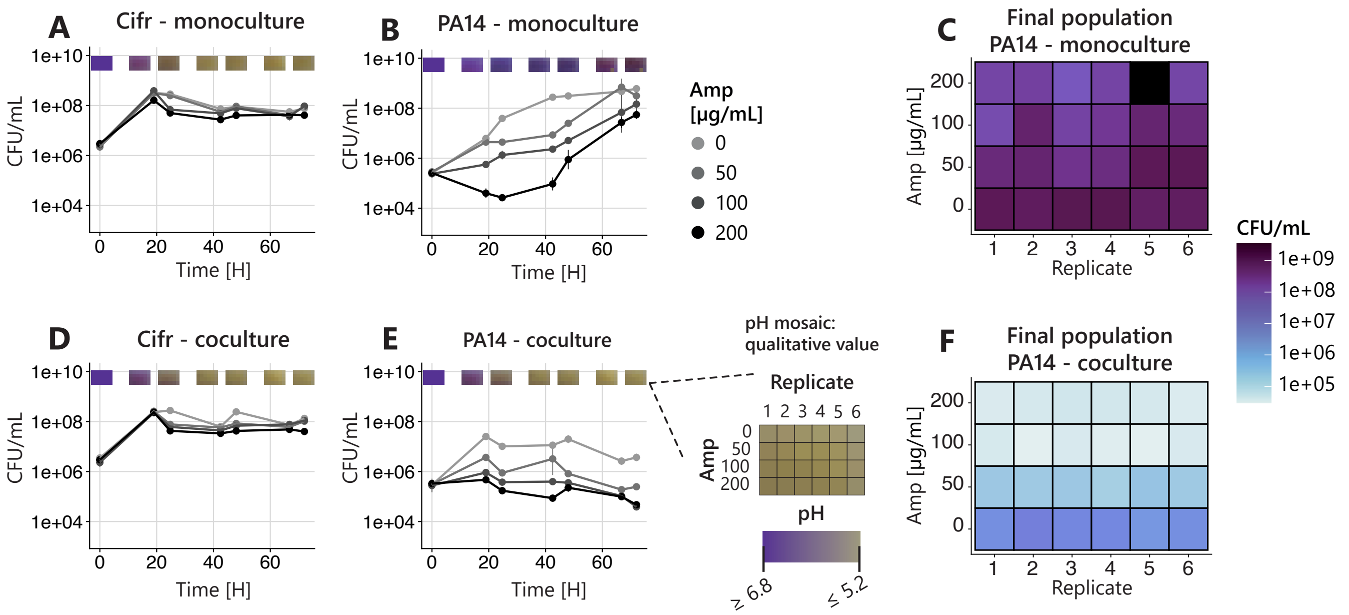


**Fig. S9.** Experiment to test the effect of frequent disturbances to the culture. *P. aeruginosa* and *C. freundii* were grown in mono- and coculture (30 mM glucose), but rather than leaving them undisturbed for 72h, CFUs were sampled twice per day (except on the day of inoculation) for 72h. pH was visually estimated using bromocresol purple and taking pictures at each sampling time. For clearer data, we performed a binary thresholding where we assigned one of 2 colors to each sample: purple for no acidification and yellow for acidification. Any culture that was in the process of turning yellow (moment of acidification) and displayed a brown/yellow color was considered as acidified, otherwise it was considered as not acidified. **(A)** Monocultures of *C. freundii*. **(B)** Monocultures of *P. aeruginosa*. **(C)** Final population of *P. aeruginosa* in monoculture from panel B. **(D)** *C. freundii* when in coculture with *P. aeruginosa*. **(E)** *P. aeruginosa* when in coculture with *C. freundii*. **(F)** Final population of *P. aeruginosa* from panel E. This experiment showed us that ampicillin had the expected inhibitory effect on *P. aeruginosa* if the cultures were disturbed and motivated us to repeat the transfer experiment again with regular disturbances.

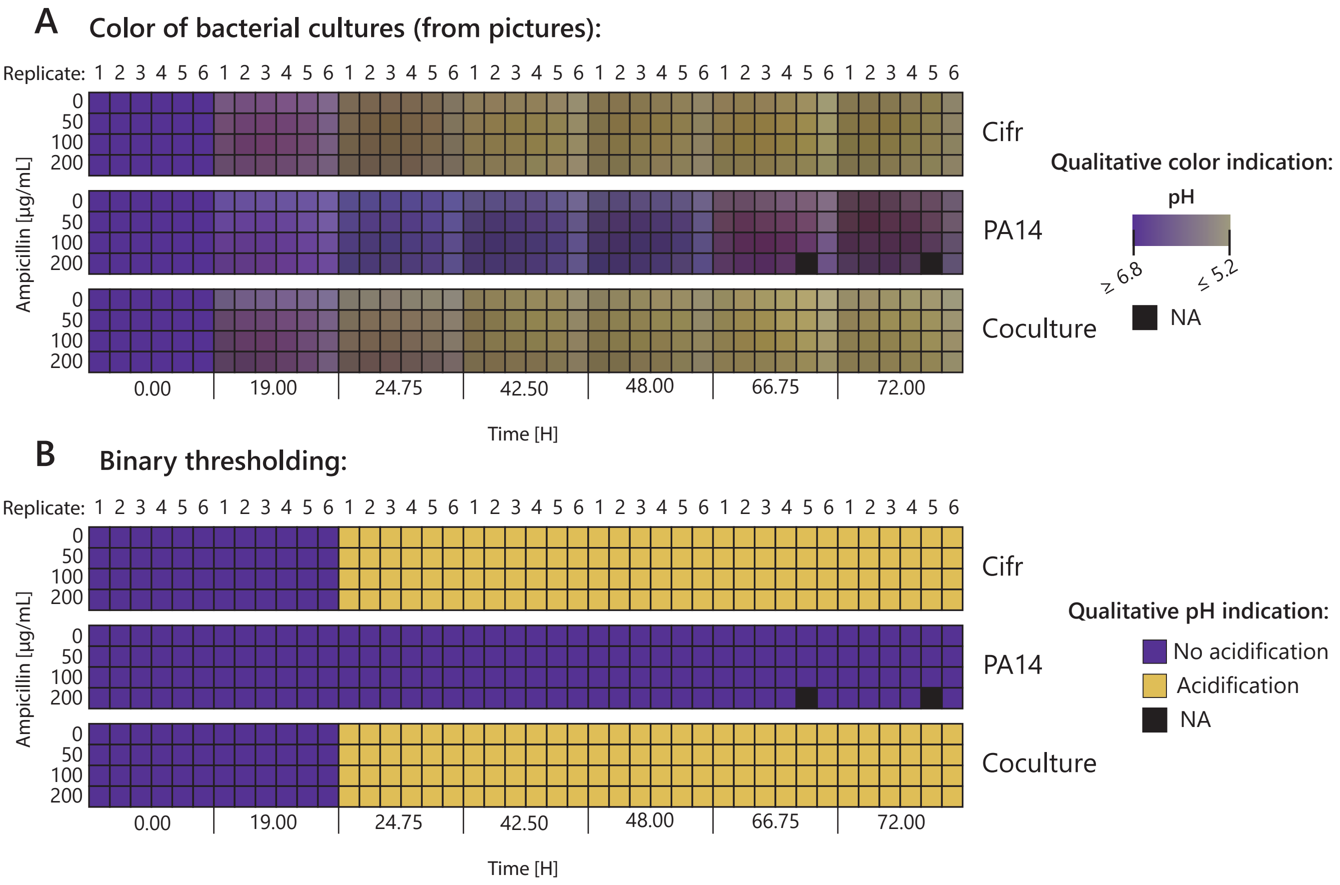


**Fig. S10.** Acidification data from Fig. S9 (A) Colors of the bacterial cultures containing bromocresol purple obtained from pictures taken before each CFU sampling. The colors of the bacteria affect the color of the culture (white/brown for *C. freundii* and pink for *P. aeruginosa* as it contains an m-Cherry tag). (B) For clearer data, we performed a binary thresholding where we manually assigned only 2 colors to each sample based on panel A: purple for no acidification and yellow for acidification. Any culture that was in the process of turning yellow (moment of acidification) and displayed a brown/yellow color was considered as acidified, otherwise it was considered as not acidified.

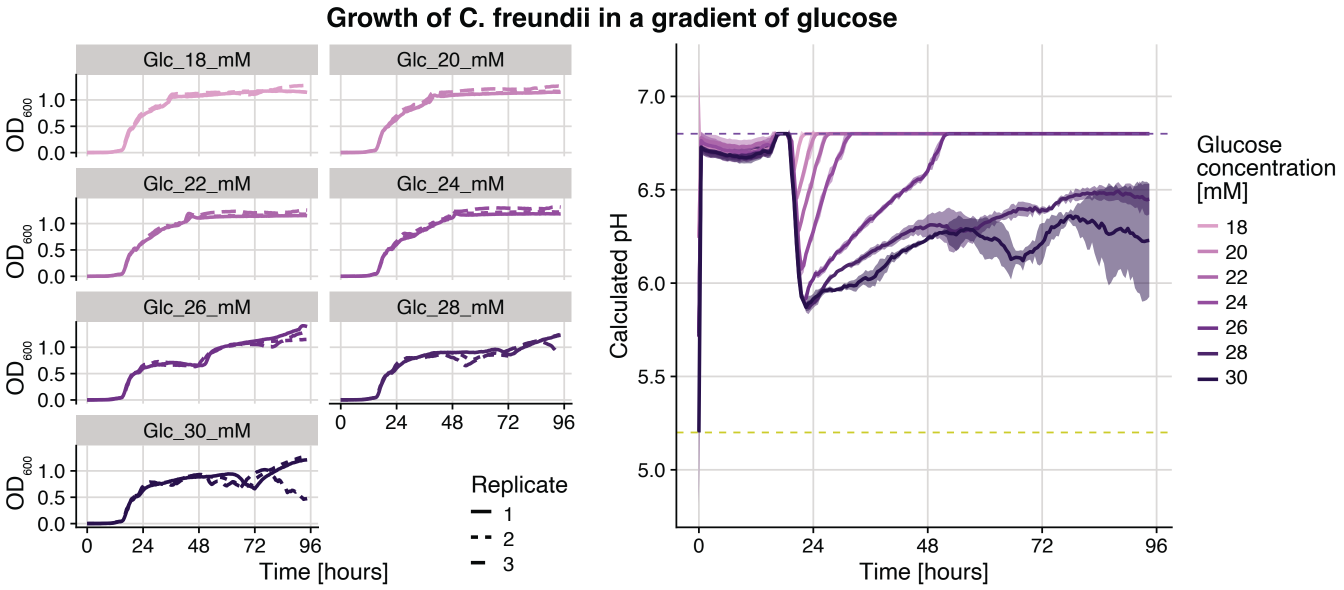


**Fig. S11.** Growth and acidification by *C. freundii* in a gradient of glucose to determine whether there was a concentration at which acidification patterns over 72h were substantially different from 15mM and from 30mM. On the right, the drop and then recovery of the pH back to 6.7 is due to the effect of the buffer (M9) in our minimal medium. This helped us to choose the concentration of 26mM for our repeated transfer experiment, where the buffer still recovers a neutral pH but the pH drops and stays low for around 24h, which we estimated would have a significant effect on *P. aeruginosa*.

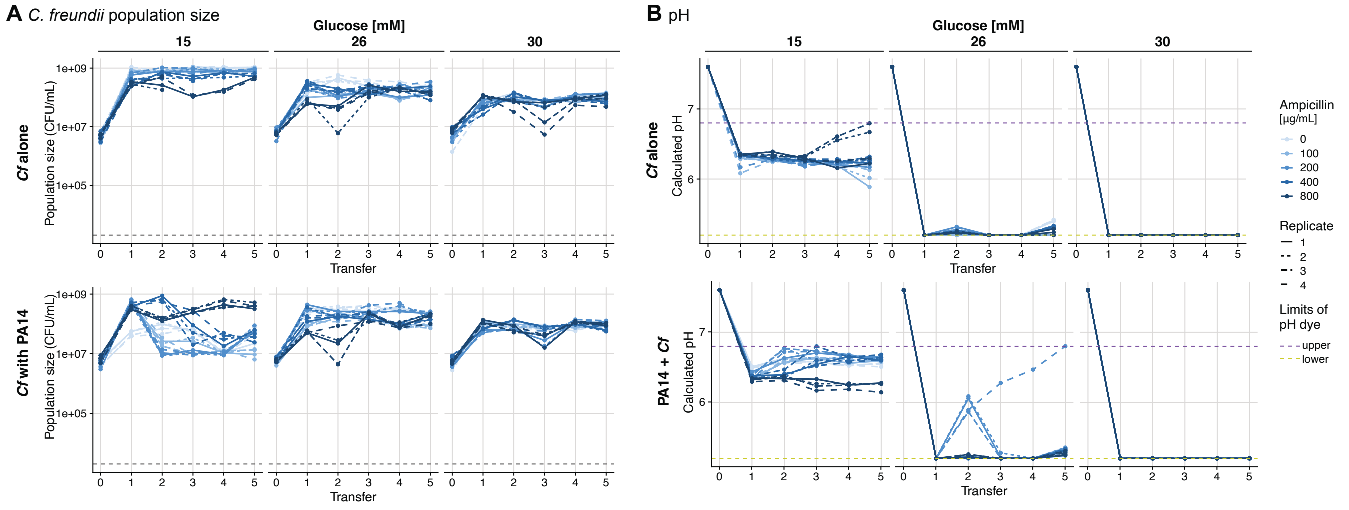


**Fig. S12.** Data for *C. freundii* from the experiment corresponding to the data shown in Fig. 3. These data show that *C. freundii* acidifies the medium more with increasing glucose concentration when alone, which has a negative effect on its population size (it suffers from its own acidification). It grows similarly at all antibiotic concentrations. In co-culture with *P. aeruginosa*, it grow less well in cases where *P. aeruginosa* grows a lot (at low glucose and low antibiotic concentration) and when it grows less, the pH also drops less.

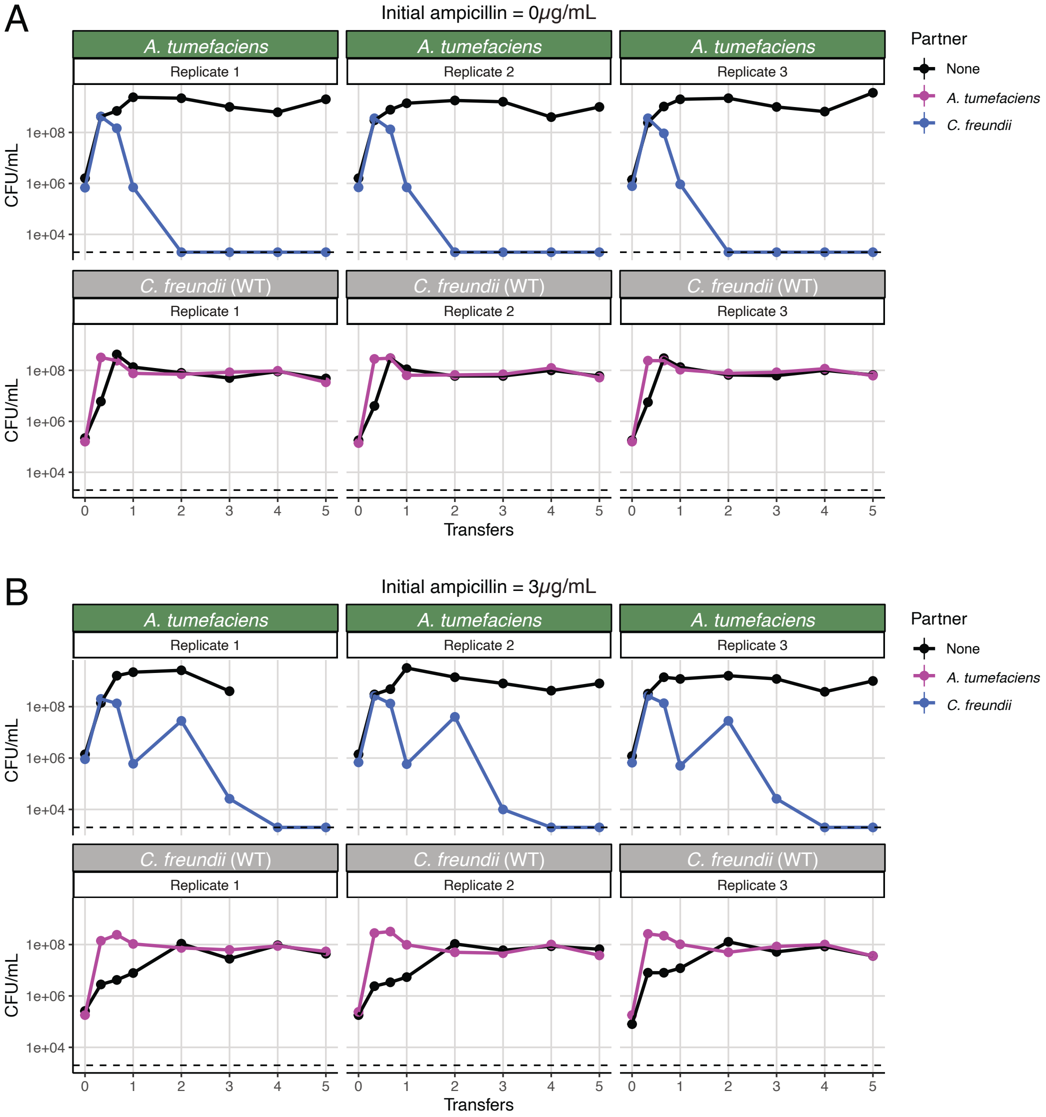


**Fig. S13.** Growth-and-dilution cycles for *C. freundii* and *Agrobacterium tumefaciens* as the pathogen. We performed 5 transfers at a 100× dilution every 72h, as in the experiment with *P. aeruginosa* (no additional disturbance). We tested two concentrations of ampicillin: 0 and 3$\mu$g/mL in panels (A) and (B) respectively. *A. tumefaciens* was eliminated by *C. freundii* at both ampicillin concentrations. It even took longer at 3$\mu$g/mL, as *C. freundii* is more sensitive to ampicillin. Note that this is because this strain of *C. freundii* is the WT, before we had selected a more resistant isolate, which is the one used with *P. aeruginosa*.

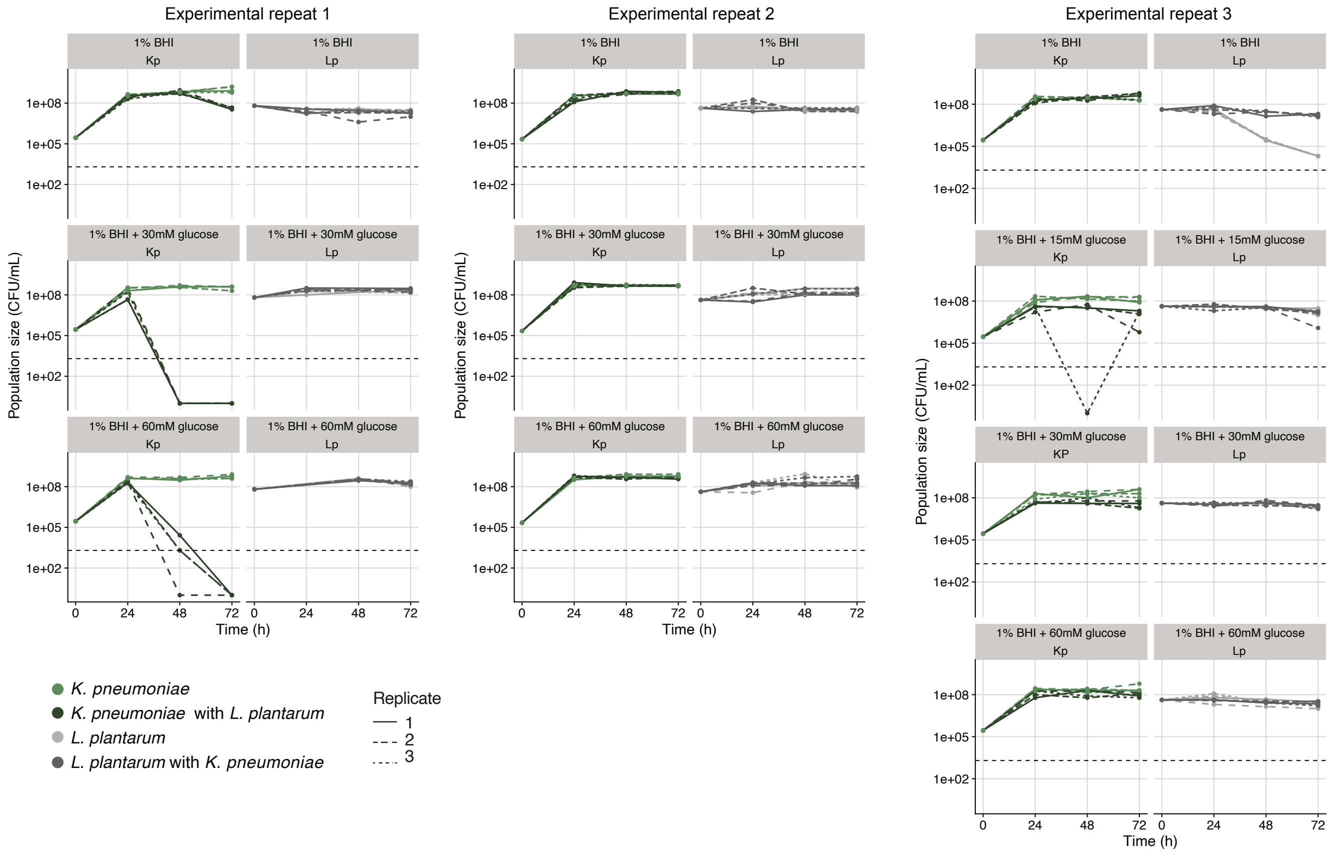


**Fig. S14.** Mono- and co-cultures (light and dark respectively) of *Lactobacillus plantarum* as the commensal and *Klebsiella pneumoniae* as the pathogen (gray and green respectively) in single batch cultures over 72 hours (no transfers). The figure shows three repeats of the experiment with different glucose concentrations shown in the rows. In the first experimental repeat, *K. pneumoniae* was eliminated at 30 and 60mM of glucose added to BHI, but not when glucose was not added. On repeating the experiment, the effect disappeared. This appears to match our observation of *K. pneumoniae*'s growth with *C. freundii*: antibiotics were needed to have a robust inhibition of the pathogen. In this case, *L. plantarum* is highly sensitive to the antibiotic, which could therefore not be used. Although we measured pH patterns with the dye, we did not add them to the figure as in all three experiments we observed no acidification without the addition of glucose, and acidification in the presence of glucose, regardless of the species composition or the concentration of glucose (as long as it was $> 0$).
